# Insect herbivory reshapes a native leaf microbiome

**DOI:** 10.1101/620716

**Authors:** Parris T Humphrey, Noah K Whiteman

## Abstract

Insect herbivory is pervasive in plant communities, but its impact on microbial plant colonizers is not well-studied in natural systems. By calibrating sequencing-based bacterial detection to absolute bacterial load, we find that the within-host abundance of most leaf microbiome (phyllosphere) taxa colonizing a native forb is amplified within leaves impacted by insect herbivory. Herbivore-associated bacterial amplification reflects community-wide compositional shifts towards lower ecological diversity, but the extent and direction of such compositional shifts can be interpreted only by quantifying absolute abundance. Experimentally eliciting anti-herbivore defenses reshaped within-host fitness ranks among *Pseudomonas* spp. field isolates and amplified a subset of putative *P. syringae* phytopathogens in a manner causally consistent with observed field-scale patterns. Herbivore damage was inversely correlated with plant reproductive success and was highly clustered across plants, which predicts tight co-clustering with putative phytopathogens across hosts. Insect herbivory may thus drive the epidemiology of plant-infecting bacteria as well as the structure of a native plant microbiome by generating variation in within-host bacterial fitness at multiple phylogenetic and spatial scales. This study emphasizes that “non-focal” biotic interactions between hosts and other organisms in their ecological settings can be crucial drivers of the population and community dynamics of host-associated microbiomes.

## Introduction

For many organisms, attack by multiple enemies is inevitable and often occurs sequentially during the lifetime of individual hosts. Prior attack can alter host phenotypes and change how future attacks unfold, often generating cascading effects at larger spatial and temporal scales ^1–4^. Given the large effects of co-infection on host health and the population dynamics of their parasites, explicitly studying co-infection is becoming increasingly common ^4–6^. But rarely has this perspective been extended to studies of diverse host-associated microbial communities (‘microbiomes’). Instead, microbiome studies tend to focus on effects of host genotype or abiotic variation on microbiome diversity patterns ^7–11^. This has left a major gap in our understanding of how host colonization from non-microbial enemies impacts the population biology of microbiome-associated taxa.

For plants, there is tremendous interest in understanding the structure and function of the microbiome both for applied purposes, such as engineering growth promotion and disease resistance ^12,13^, and as model systems for host–microbial symbioses more generally. Insect herbivory represents a pervasive threat to plants in both native and agricultural settings ^14^. Herbivory alters plant phenotypes through tissue damage and induction of plant defenses, which can change susceptibility of plants to attack by insects ^15^ as well as microbes ^16,17^. Thus, factors that influence the impact of herbivores on hosts will likely affect the colonization and growth of plant-associated microbes. While insect herbivores ^14^ and plant-associated microbes have clear effects on plant phenotypes and fitness ^18^, they are generally considered independently. Our study addresses this gap by explicitly considering how patterns of abundance and diversity of leaf-colonizing (endophytic) bacterial taxa are altered in the presence of insect herbivory and by exploring the associations among herbivory, bacterial infection, and plant fitness in a native forb (*Cardamine cordifolia*, Brassicaceae; ‘bittercress’).

We first used marker gene sequencing (16S rRNA) coupled with paired leaf culturing to establish and validate sequence-based estimates of absolute bacterial load in host tissue. By elucidating a relationship between bacterial load and the sequence counts of bacteria-versus host-derived 16S (**Box 1**), our approach enabled standard 16S marker gene sequencing to quantitatively reveal variation in abundance distributions of entire suites of bacterial taxa across hosts with and without prior insect herbivory. We then assessed the extent of co-clustering between microbial abundance and intensity of insect herbivory at the plant patch scale across our study populations, and we related microbe–herbivore co-aggregation to fruit set, a component of plant fitness. In parallel, we directly examined variation in sensitivity to inducible plant defenses among 12 genetically diverse, bittercress-derived isolates of *Pseudomonas* spp. bacteria. We did so by experimentally infecting accessions of native bittercress in the laboratory in which prior herbivory was simulated by exogenously pre-treating plants with the plant defense hormone jasmonic acid (JA).

Our experiments reveal that insect herbivory, via induction of plant defenses, can modify endophytic bacterial diversity patterns by amplifying naturally prevalent and potentially phytopathogenic bacterial taxa within a native plant host. This mechanism may be at least partly responsible for the strong positive association between herbivory and endophytic bacterial abundance within leaf microbiomes seen under field conditions. Crucially, the patterns and degree of bacterial abundance variation we found cannot be revealed by traditional compositional analysis of high-throughput marker gene sequencing, which masks the extent and direction of within-host variation in bacterial load. By linking marker gene counts to an absolute standard, our study reveals how insect herbivory associates with variation in bacterial loads at leaf and patch scales within a natural plant population. More generally, this work highlights the importance of (*a*) accounting for prevalent but “non-focal” biotic interactions hosts have with other colonizers in their natural contexts, and (*b*) using detection and analytical approaches to quantify these effects on components of microbial fitness.

## Methods

### Field studies of herbivore–bacteria co-infection

We surveyed herbivore damage arising from the specialist leaf-mining insect *Scaptomyza nigrita* (Drosophilidae; Fig. S1f–g) in replicate 0.5 m^2^ plots of native bittercress along transects in sub-alpine and alpine streams near the Rocky Mountain Biological Laboratory (RMBL) in each of two years (2012, Emerald Lake, EL, *n* = 24 plots; 2013, site North Pole Basin, NP, *n* = 60 plots; Fig. S1a–d). Although not our primary focus, these field studies were also designed to test how mid-season pre-treatment with the exogenous plant defense hormones JA or salicylic acid (SA) impacts plant attack rates by *S. nigrita*. JA induces canonical defenses against chewing herbivores in plants ^17^, including bittercress, and JA-induced bittercress can locally deter adult *S. nigrita* and reduce larval feeding rates ^19^. SA treatment canonically induces defenses against biotrophic microbial colonizers and often pleiotropically suppresses plant defenses against chewing herbivores ^17^, including *S. nigrita* ^20^. Thus, treatment with either plant defense hormone has the potential to modify the foraging behavior of S. nigrita. Our analysis of the impact of hormone treatments on *S. nigrita* foraging patterns was previously published for site EL ^20^, and we implemented a similar approach for site NP in this study. Full experimental design details are given in the Supplemental Methods and are depicted in the schematic in Fig. S1e.

By the end of the growing season, when herbivory and bacterial infection had run their course, we determined *S. nigrita* leaf-miner damage status of all leaves (both sites) as well as fruit set (site NP only) produced on each of the focal bittercress stems (stem-level sample size *n* = 768, site EL; *n* = 1920, site NP). At both sites, we collected leaf tissue in a randomized manner (see Supplemental Methods) to quantify the abundance and diversity of bacteria that had colonized the leaf interior.

### Amplicon sequencing of bacteria in leaf tissues

We quantified bacterial abundance in leaf tissues using next-generation amplicon sequencing of the bacterial 16S rRNA locus using the Illumina MiSeq platform. In order to enrich our samples for endophytic bacteria, we surface-sterilized all samples prior to DNA extraction, which achieved a reliable reduction of bacterial abundance as detected by our 16S analysis approach (Fig. S2). Subsequently, we extracted DNA from the 192 leaf discs (∼ 0.8 cm^2^) from site EL and 192 tissue pools from site NP (4 four discs per pool) and amplified bacterial 16S following protocol established for the Earth Microbiome Project (see Supplemental Methods). We amended this protocol by including peptic nucleic acid (PNA) PCR clamps into reaction mixtures to reduce amplification of host chloroplast- and mitochondria-derived 16S, following Lundberg et al. ^21^. This was highly effective at reducing the proportion of host-derived 16S reads per library in our sample sets (Fig. S3).

We then used `DADA2`^22^ to error-correct, trim, quality-filter, and merge the paired-end sequencing reads that passed error thresholds off the sequencer. Of the approximately 4 million raw reads, ∼ 90% were retained following quality control via `DADA2` (Fig. S4), and these reads were then delineated into exact amplicon sequencing variants (ASVs). 16S reads from bittercress chloroplast or mitochondria were manually curated and summed into ‘host-derived’ for comparison with bacteria-derived 16S (see Supplemental Methods).

### Quantifying and modeling bacterial abundance patterns

In order to quantitatively asses how herbivore damage relates to abundance and diversity of microbial plant colonizers, we required a link between 16S counts and bacterial load. We therefore devised and validated an estimator (*γ*) of the abundance of bacterial ASVs within host tissues (**Box 1**). Using *γ* as an empirically validated estimator of absolute bacterial load in leaf tissues, we then constructed a two-stage modeling approach to estimate bacterial load across our complete sample set.

We first fit and compared a series of increasingly flexible Bayesian regression models to estimate how *γ* varies as a function of herbivore damage in leaves (see Supplemental Methods). When calculating *γ*, we took *r*_*B*_ at the bASV-level for all bASVs within each of the 14 most abundant bacterial families, together comprising > 95% of total bASV counts in the datasets. We then took the candidate best stage-1 model, heuristically defined as the model with the lowest leave-one-out Bayesian information criterion (LOO-IC) ^23^, and used it to generate *n* = 200 replicate sets of simulated response values 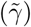 predicted by the model parameters fit to the original data.

In the next stage, we used this distribution of 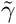 as an input predictor variable to the model we fit between our observed *γ* and observed log CFU (**Box 1**). This allowed us to report bacterial abundance estimates, based initially on 16S count data, on the scale of predicted log CFU per unit leaf mass—a more directly interpretable measure of within-host fitness. Rather than point estimates, we sampled intercept, slope, and residual error parameters from their joint posterior distribution of the calibration CFU model for each data point independently. Specifically, this has the effect of incorporating noise in the fit between observed log CFU and *γ* such that downstream predictions are not overly biased by the precise value of any regression slope estimate, which may itself arise from peculiarities in the action of PNA during amplification of 16S. Overall, this two-stage modeling approach was designed to incorporate uncertainty in the model fit for *γ* as well as in the relationship between observed *γ* and observed log CFU. The endpoint of this approach is 200 sets of posterior predicted log CFU values for each sample in the dataset, which formed the basis of downstream calculations of bacterial abundance variation, as well as ecological diversity (Shanon evenness *J*′) and similarity (Shannon–Jensen divergence *SJ*) in and between damaged and undamaged leaf sets, respectively (see Supplemental Methods).

### Population-level analysis of herbivore–bacteria co-aggregation

We assessed how patch-level variation in herbivory correlated with bacterial infection intensity at the field-scale by focusing on the most highly abundant bASV at both field sites (‘`Pseudomonas3`’). Leaf-level abundance of this individual bASV was predicted using the approach described above. We then summed the predicted abundance (on the linear scale) of bacteria across leaves within each plant patch and used these predicted bacterial sums to calculate the cumulative proportion of the total `Pseudomonas3` population harbored by plant patches with differing levels of herbivory, which portrays the extent of co-aggregation of herbivores and bacteria across the host population.

### Experimental infections *in planta*

We directly examined how inducible plant defenses against chewing herbivores impacted within-host bacterial performance using field-derived accessions of bittercress plants and their bacterial colonizers. We randomized the selection of six focal strains from within each of the two dominant *Pseudomonas* clades (*P. syringae*, and *P. fluorescens*) represented in our endophytic strain collection from bittercress ^20^. We infected each strain into a single leaf on each of *n* = 32 distinct bittercress clones that had been randomized to receive pre-treated 3 days prior with JA (1 mM; Sigma) or a mock solution. Bittercress clones were originally isolated as rhizomes from various sites within 2 km of the RMBL in 2012 and were re-grown in the greenhouse at University of Arizona for up to 12 months prior to use ^24^. Two days post infection, we sampled, sterilized, homogenized, and dilution-plated leaf discs onto non-selective rich King’s B media, following Humphrey et al. ^20^. We compared bacterial abundance (log _2_) between treated and untreated samples using Gaussian Bayesian regression models. We subsequently used posterior predicted abundances as the basis for considering how herbivore-inducible defenses impact the composition and diversity of this *Pseudomonas* community at different taxonomic levels (see Supplemental Methods).

## Results

### Bacterial loads are amplified in insect-damaged leaf tissues

By linking information from 16S counts to absolute bacterial abundance (**Box 1**), we found that bacteria detected within herbivore-damaged leaves exhibit local population sizes several doublings greater compared to the bacteria found in undamaged leaves (Fig. 1a,b; median ± 95% credible interval of posterior predicted additional doublings: 2.5 [2.1; 3.9] site EL; 4.5 [3.6; 5.3] site NP). This result, rooted in sequence data, is further validated by the parallel observation that damaged leaves showed higher bacterial loads than undamaged leaves via culturing of the *n* = 101 calibration set (Fig. S5; mean difference of 3.7 bacterial doublings [1.8–5.6, 95% c.i. on mean difference]; Welch’s unequal variance two-sample *t*-test, *t* = 3.86, *p* < 0.001), which is quantitatively consistent with a prior result from a parallel and independent culture-based study in this system ^20^.

**Figure 1:**
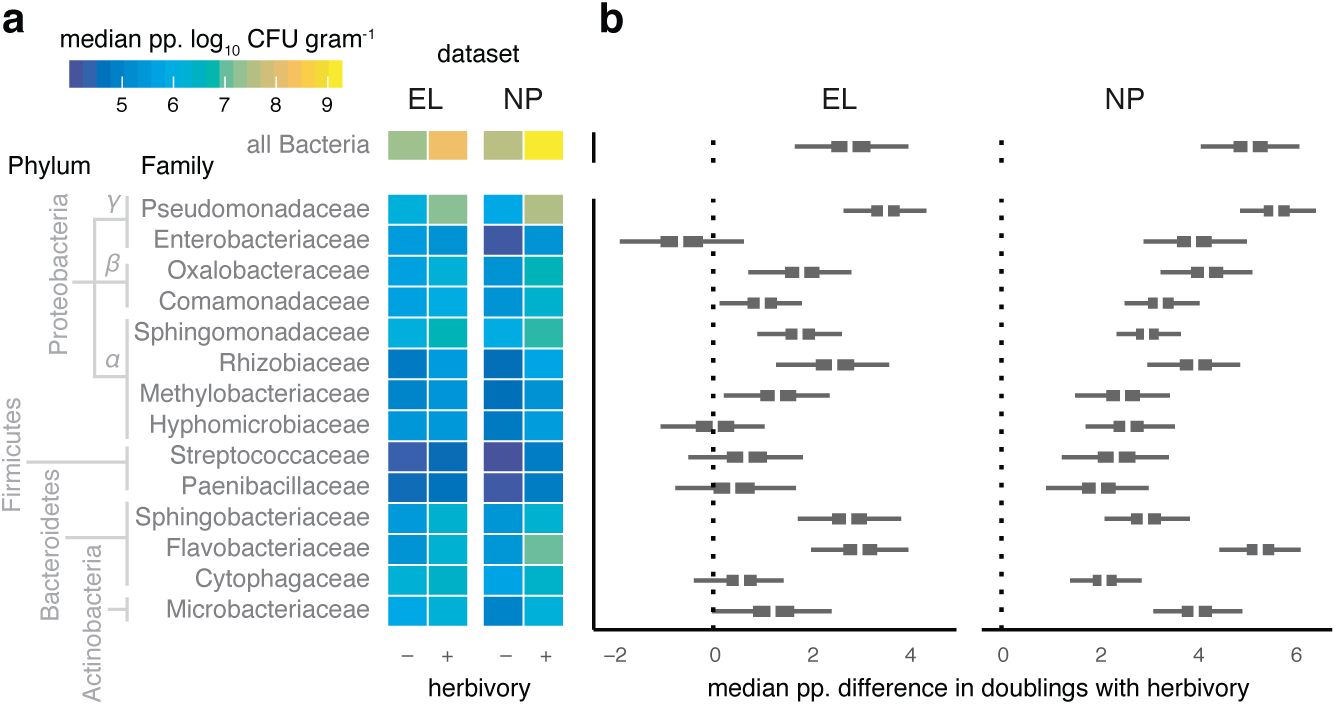
Pervasive increases in endophytic bacterial load in herbivore-damaged leaves. **a–b**. Posterior predicted (‘pp.’) infection intensity of bacterial amplicon sequence variants (bASV) from the 14 most prevalence bacterial families show variation in the extent of elevated growth in herbivore-damaged leaf tissue. **a**. Heatmap shows median predicted log _10_ bacterial abundance (colony-forming units, CFU) per g starting leaf material) from 200 posterior simulations of the best-fitting model of each bacterial family separately (see Methods). **b**. Median (white), 95%, and 50% quantiles of the median difference in the number of predicted bacterial cell divisions (i.e., doublings) achieved in herbivore-damaged leaves compared to undamaged leaves, for sites EL and NP separately.

### Herbivore-associated bacterial amplification is both community-wide and taxon-specific

We then capitalized on the high taxonomic resolution and sampling depth afforded by amplicon sequencing to examine shifts in abundance and distribution of diverse bacteria within the bittercress leaf microbiome. Within-host density of bacterial bASVs from several bacterial families was elevated in herbivore-damaged leaves compared to undamaged leaves (Fig. 1). For most families, the relative increase in within-host density with herbivory was greater at site NP than at site EL (Fig. 1b), but this was largely because several taxa showed lower baseline loads in undamaged leaves at site NP compared to site EL (Fig. S6a). In contrast, bacterial loads for all families were similar for damaged leaves at both site (Fig. S6a,b). Pseudomonadaceae was the most abundant taxon across all leaves and also showed the greatest fold increase under herbivory (Fig. 1).

Several family-level *γ* models showed support for bASV-level differences in intercept and slope values (Table S2), including Pseudomonadaceae and Sphingomonadaceae. Two individual bASVs in particular drove family-level patterns in these clades (Fig. S7), which together comprised ∼ 20% of all sequencing reads across the two sample sets. Within the Pseudomonadaceae, `Pseudomonas3` was the most abundant bASV, which falls within the putatively phytopathogenic *P. syringae* clade (Fig. S8). We previously showed that *P. syringae* strains can be pathogenic, induce chlorosis, and reduce leaf photosynthetic function in bittercress ^20^. Thus, a major component of the signal of elevated bacterial load in the presence of insect herbivory comes from putatively phyopathogenic genotypes within the group *P. syringae*.

### Compositional shifts in leaf bacterial communities under herbivory

When the absolute bacterial abundance patterns described above were analyzed in a compositional framework, we detected differences in overall community structure and ecological diversity between damaged and undamaged leaves. Specifically, we found lower evenness (*J*′; Fig. 2b) in damaged leaves, indicating a stronger skew towards a smaller number of bacterial taxa: Pseudomonadaceae comprise an even greater proportion of the population in damaged leaves owing to their already-high average abundance in undamaged leaves and large fold-increase under herbivory. Family-level relative abundances differed in terms of Shannon–Jensen divergence (i.e., *β*-diversity) between damaged versus un-damaged leaves (Fig. 2c), indicating that amplification of bacteria in herbivore-damaged leaves can produce community-wide signatures of reduced within-host diversity and elevated between-host diversity at broad taxonomic scales.

**Figure 2:**
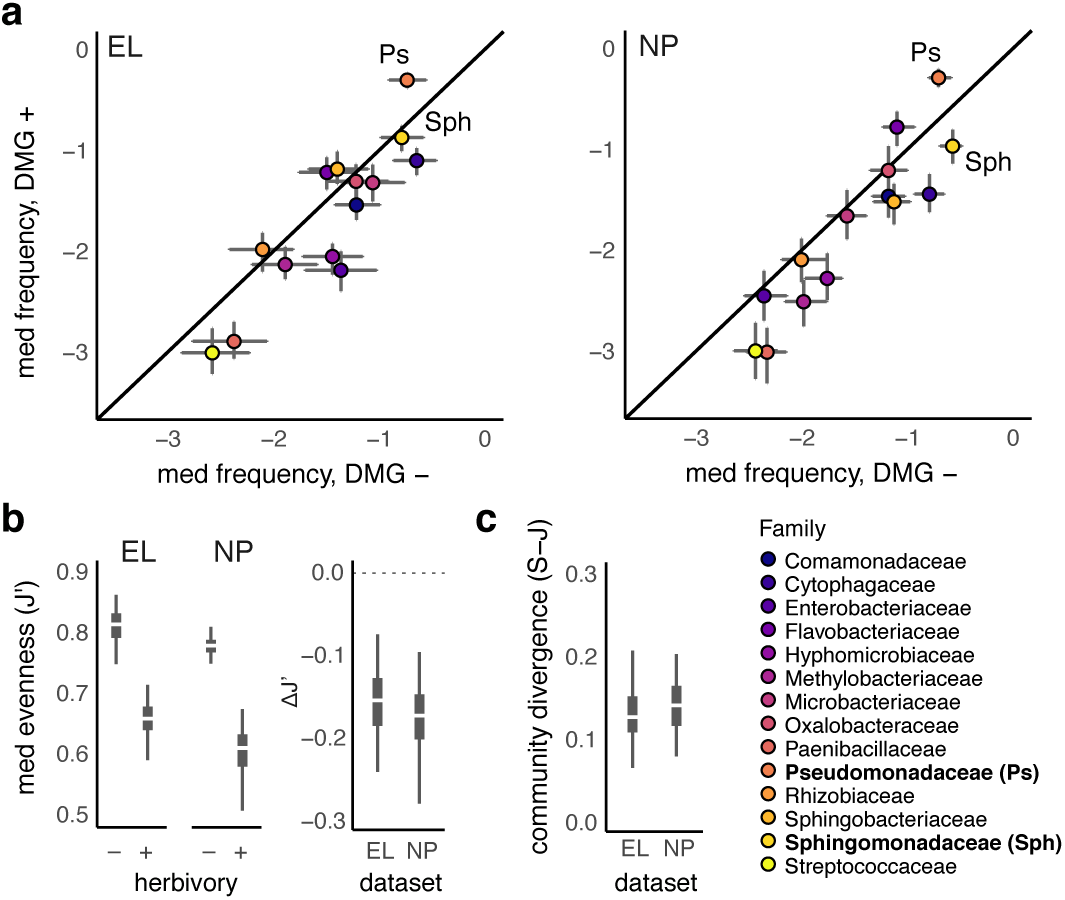
Herbivore-damaged leaves harbor compositionally diverged microbiomes with reduced ecological diversity shifted heavily towards Pseudomonadaceae. **a**. log _10_ relative abundance of each family in undamaged (*x*-axis) versus damaged (*y*-axis) samples shows skew towards Pseudomonadaceae (Ps) and relative reductions of abundance among most other taxa at both study sites, including *Sphingomonas* (Sph) which shows a ∼ 2-fold increase in number of doublings in herbivore damaged leaves. **b**. Compositional changes from the amplification of already abundant taxa (e.g., Pseudomonadaceae) produces reduced community-level evenness (*J*′) and leads to compositional divergence (i.e., *β*-diversity) between damaged and undamaged leaves at both study sites (**c**).

### Plant defenses against chewing herbivores enhance growth of putative phytopathogens *in planta*

Plant pre-treatment with the plant defense hormone JA caused statistically clear alterations in within-host growth of five of the twelve *Pseuomonas* spp. strains tested (Fig. 3a), with the most pronounced changes resulting in 2.5–5 additional doublings of two phylogenetically distinct *P. syringae* isolates (20A and 02A; Table S3). Increased within-host density of these two strains can account for differences in total *Pseudomonas* abundance, as well as differences in abundance patterns summed at the level of bacterial clade (*P. syringae* versus *P. fluorescens*; Fig. 3b). By recapitulating the elevated *P. syringae* and family-wide increased abundance under herbivory seen in our field studies, this greenhouse result highlights that induction of plant defenses against chewing herbivores is one potential mechanism whereby insect herbivory could lead to amplification of bacterial taxa within the bittercress leaf microbiome.

**Figure 3:**
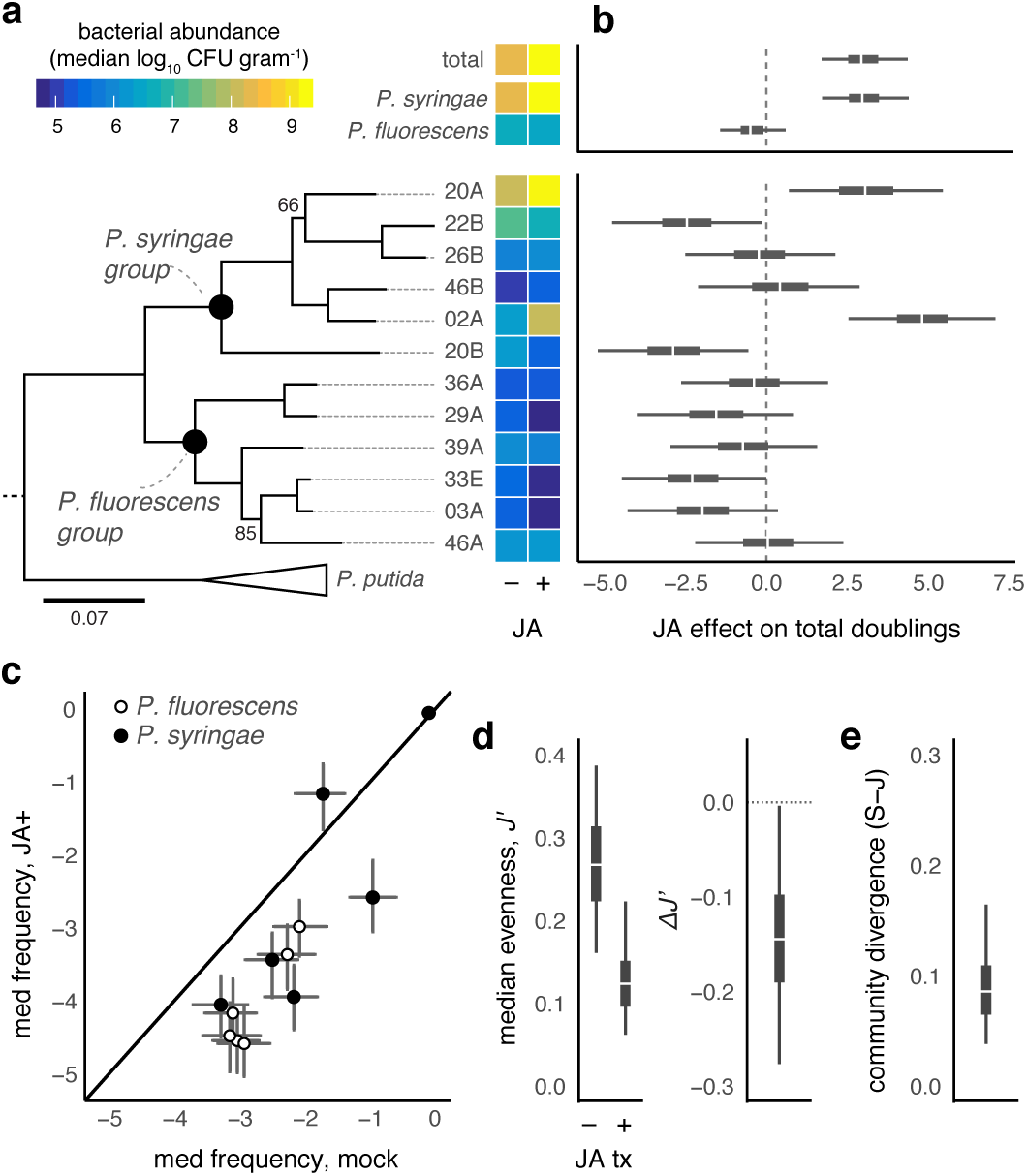
Eliciting plant defenses against chewing herbivores alters within-host performance of putative phytopathogens. **a**. Experimental infections with 12 *Pseudomonas* spp. strains, concurrently isolated from EL study site ^20^, show strain-to-strain variation in growth under mock-treated (M) and jasmonic acid (JA) induced plants. Heatmap shows median log _10_CFU g^−1^ surface sterilized plant tissue 2 d post inoculation. Maximum likelihood phylogeny of strains estimated with four housekeeping loci (2951 bp) from ^20^. **b**. Median, 95%, and 50% quantiles of the posterior difference between the number of bacterial doublings attained by bacteria growing in JA-versus mock-treated leaves (see Supplemental Methods). **c**. Compositional analysis of relative abundances calculated from (**a**) reflect decreased evenness (*J*′; **d**) in JA-treated plant tissues, leading to overall community-level divergence (**e**). Median, 95%, and 50% quantiles from 200 posterior simulations of abundance (**c–e**) from the best-fitting model fit to data in (**b**).

Notably, two strains (22B and 20B) exhibited markedly decreased within-host fitness in JA-treated compared to mock-treated leaves (Fig. 3a–b). Such herbivore-driven fitness variation among *P. syringae* is undetectable when only considering larger taxonomic scales of genus or family (Fig. 3), where genotypes which *increase* in local abundance contribute to an overall signature of elevated taxon-wide abundance as measured by lower resolution tools (e.g., 16S sequencing). Thus, induction of plant defenses against chewing herbivores leads to the amplification and numerical dominance of a narrow subset of the *P. syringae* community within this host population (Fig. 3c). Such changes result in compositional shifts towards decreased Shannon evenness in JA-treated leaves (Fig. 3d) and an overall community-wide divergence with mock-treated leaves (Fig. 3e).

### Putative phytopathogens are aggregated in highly herbivore-damaged plant patches

We then analyzed how `Pseudomonas3`, a highly abundant individual bASV within the *P. syringae* group, varied across bittercress plant patches in relation to the level of herbivory on those plant patches. At site NP, we found a highly aggregated (i.e., right-skewed) distribution of herbivore loads across plant patches (Fig. 4a, top marginal density plot). This aggregated herbivore distribution across plant patches results in a predicted 50-fold enrichment of local density of `Pseudomonas3` in the most-damaged compared to the least-damaged plant patch (Fig. 4). Analyzed in a more general framework, over half the predicted `Pseudomonas3` population is harbored in just one fifth of the plant patches in the bittercress population at our study site NP (Fig. S9).

**Figure 4:**
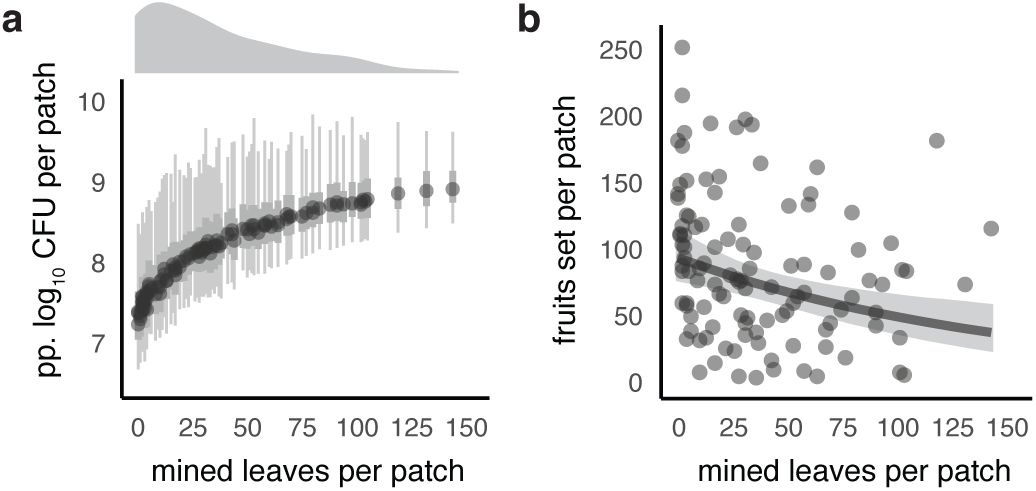
Co-infection by herbivores and phytopathogenic bacteria is aggregated across plant populations and is associated with lower plant reproduction. **a**. Median, 95%, and 50% quantiles of 200 posterior simulations of predicted (‘pp.’) bacterial load across plant patches (*n* = 110 at site NP; *n* = 16 stems sampled per patch). Density plot above *x*-axes exhibits right-skewed (i.e., aggregated) distribution of herbivore damage at the plant patch level. **b**. Patch-level herbivory (and thus co-infection intensity) is associated with decreased fruit-set in this native plant population. Plotted are raw fruit-set data summed at the patch level (*n* = 16 stems per patch), with marginal effects slope (and its 95% credible interval) plotted after accounting for average plant height (see Table S4).

### Herbivore–bacteria co-aggregation is associated with lower plant fitness

At site NP, bittercress patches with higher herbivore intensity showed lower reproductive success, with the most damaged patches estimated to produce half as many fruits as patches with no herbivore damage (Fig. 4b, Table S4). Plants with more insect damage tend to have higher levels of bacterial infection. Thus, standing variation in fruit set is closely associated with levels of co-aggregation of these plant natural enemies across our sample within this native bittercress population.

### Causes of herbivore aggregation in natural plant populations

The degree of herbivore aggregation among host plants at site NP varied extensively across plant patches at site NP (Fig. S10a–b); the marginal effects on herbivore damage arising from early-season treatments with plant defense hormones SA or JA were small (Fig. S10c). Estimates for both SA and JA treatment coefficients were elevated above the mock/control condition, but the posterior distribution for both hormone effects overlapped zero (lower 4th %-ile < 0 for JA; lower 15th %-ile < 0 for SA; Table S5). Thus, while prior plant exposure to JA, and possible also SA, may cause elevated *S. nigrita* herbivory at the patch scale, standing variation in *S. nigrita* herbivory arising stochastically or from unmeasured factors at site NP dominates over any causal effects of our population-level defense hormone manipulation. Additionally, these plant-level defense hormone treatments (five weeks prior to leaf sampling) showed no discernible effect on distributions of *γ* for overall or family-wise bacterial abundance (Fig. S11).

## Discussion

### Overview

Here we show that insect herbivory is strongly associated with bacterial abundance and diversity within a native plant microbiome using field and greenhouse experiments. We provide evidence that activation of plant defenses against chewing herbivores is at least one causal mechanism whereby such within-host amplification of leaf-colonizing bacteria can occur. Specifically, the growth of a majority of bacterial taxa found in the leaf microbiome of native bittercress was amplified in plant tissues damaged by the specialist herbivore *S. nigrita* (Fig. 1) at two separate sub-alpine field sites. These ecological effects were only detectable by linking sequence-based bacterial quantification to an external standard of absolute abundance (**Box 1**), rather than relying on compositional analysis as is commonly done with studies of both plant and animal microbiomes (but see Vandeputte et al. ^25^).

The bacterial clades most altered under herbivory include strains from groups well-known for causing plant disease (*P. syringae*), and our follow-up experimental work in the greenhouse showed that inducing plant defenses against chewing herbivores in bittercress was sufficient to cause similar degrees of amplification of putatively phytopathogenic *P. syringae* genotypes in leaf tissues. Amplification of specific *P. syringae* genotypes can largely account for species- and family-level patterns seen in our field studies, which has coarser taxonomic resolution. Overall, these experiments suggest that *S. nigrita* herbivores may play a larger role than previously appreciated in promoting the within-host growth of particular bacterial genotypes or pathovars in bittercress, although the causal nature of the role of herbivores was not gleaned from the field portion of this study. Given that the majority of plants face herbivore attack to some degree ^14,26^, it is possible that our results generalize across plant–microbe systems.

### Herbivore-inducible plant defenses can amplify putative phytopathogens

The mechanisms governing growth-promoting effects of insect herbivory on leaf-colonizing bacteria are potentially numerous. Leaf damage itself can release nutrients, alleviating resource constraints for bacteria while also creating routes for colonization of the leaf interior from the leaf surface ^27,28^. Plant defenses induced by chewing herbivores could directly or indirectly alter interactions with bacteria independent of the physical effects of plant tissue damage. It is known that JA-dependent anti-herbivore defenses can suppress the subsequent activity of signaling pathways responsive to bacterial infection ^17^, allowing bacteria (including strains of *P. syringae*) to reach higher densities within JA-affected leaf tissues ^29^. Insects such as *S. nigrita* trigger such JA-dependent host defenses in bittercress ^19^, and this form of defense signaling is widely conserved among diverse plant groups ^17^. Released from top-down control, a diversity of resident microbes can then proliferate as defenses are more strongly directed against herbivory, which may manifest in community-wide patterns of abundance changes as noted in our study (Fig. 1).

The hypothesis that anti-herbivore defenses pleiotropically increase bacterial growth is consistent with results from our greenhouse experiment. We found that the within-host fitness of several strains of putatively phytopathogenic *P. syringae* increased within bittercress leaves pre-treated with JA compared to mock-treated leaves (Fig. 2). Although our experiment is consistent with this plausible mechanism by which herbivores can facilitate bacterial growth within plants, it does not identify the proximal mechanism(s) responsible for these effects. JA-induced defenses may have instead stressed the plants, decreasing basal tolerance to infection, or shunted resources towards investments which reduce the ability of plants to resist or tolerate bacterial infection ^30^. Such net effects of JA induction on bacterial abundance are likely contingent on underlying constitutive levels of genetic resistance and/or tolerance to herbivory, traits which often exhibit quantitative variation within and among plant populations ^31^. The role of host genetic variation in defense responsiveness phenotypes in mediating the impacts of herbivore attack on microbial plant colonization is an open avenue of future research.

Finally, several other abundant bacterial groups (e.g., Sphingomonadaceae, Flavobacteriaceae) exhibited amplified abundance under insect herbivory in our field studies (Fig. 1). Functional studies examining finer-scale variation among genotypes of these relatively less well-studied bacterial groups would be highly fruitful for establishing a more general understanding of the mechanistic basis of plant–microbe interactions in the context of inducible defenses against chewing herbivores.

### Herbivore distributions can alter the spatial patterning of plant disease

The impact of insect herbivory on phyllosphere bacteria can be observed at several spatial scales. Herbivore damage was highly clustered on a subset of hosts (Fig. 4), which is a pattern consistent with other plant–herbivore systems ^32^ as well as many host–macroparasite systems more generally ^33^. A potential accompanying effect of aggregated herbivore damage is to enrich bacterial infection on a subset of the host population (Fig. 4, Fig. S8), altering the spatial structure of growth and, potentially, transmission of plant-colonizing microbes. Uncovering the temporal dynamics of how herbivore aggregation precedes or follows microbial attack—or whether the two colonizers cyclically amplify one another—will require more controlled studies that manipulate the timing and density of herbivory itself.

Regardless of the precise mechanisms resulting in such herbivore–microbe co-aggregation, plant patches with the highest levels of co-aggregation had substantially (∼ 50%) lower reproductive success compared to minimally-damaged plant patches (Fig. 4b). Although we cannot identify the cause of lower plant fitness from our study, the co-aggregation of these distinct plant colonizers may at least partly explain it. Whether these plants were more stressed to begin with or achieved lower fruiting success because of their infestation with herbivores and phytopathogens cannot be resolved without future studies which isolate the causal effects of single and multiple infection on plant fitness.

What drives the highly skewed pattern of herbivory among host plants? Plant populations often display a patchwork of defensive phenotypes, influenced by plant abiotic stress, variation in the underlying defensive alleles, or by defense induction from prior herbivore or microbial attack. While plant defenses can shape herbivore attack rates in the laboratory and over wide spatial extents ^31^, less is known about how patterns of defense induction impact the population dynamics of insect herbivores ^32,34^. Many insects are deterred by anti-herbivore defenses, but some specialist herbivores use these same cues as attractants owing to detoxification mechanisms which confer resistance to such defenses ^35^. *S. nigrita* uses anti-herbivore plant defenses to locate host but also avoids high concentrations of defensive compounds when given the option ^19^. Thus, the joint expression of positive chemo-taxis towards JA-inducible compounds, coupled to aversion of high levels of JA responsive defensive chemistry, may influence where herbivory becomes concentrated among plants in native bittercress populations.

Results from our field hormone treatments using JA and SA both showed elevated patch-level *S. nigrita* leaf miner damage compared to mock-treated patches (Fig. S10). However, the statistical signature of these treatment effects was not clear enough to confidently conclude that our field trials substantially altered natural patterns of herbivory by this specialist, given the high degree of stochastic or unexplained variation in herbivore damage we observed across plant patches (Fig. S10, Table S5). Discovering the biotic and abiotic factors structuring herbivory patterns in natural host populations thus remains a challenge in this system ^36^ as well as many others ^34^, and we have not attempted to solve this problem in the present study. Nonetheless, our study suggests that predicting herbivore distributions may be key for understanding population-level distributions of plant-associated bacteria. Regardless of its causes, insect herbivore damage can be readily measured and incorporated into plant microbiome studies in order to help reveal the drivers of variation in microbiome abundance and diversity within plant communities.

### Quantifying bacterial loads is crucial for understanding the ecology of the microbiome

The patterns of abundance variation among bacterial taxa across leaf types, when distilled into a community-level compositional metric, showed decreased ecological diversity (i.e., evenness) in damaged versus un-damaged leaves, resulting in overall compositional divergence between sample sets (Fig. 2–3). This results from particular taxa undergoing larger absolute changes in abundance than other taxa, which leads to stronger skews in the composition of the community calculated on the relative scale. Compositional analysis on its own would preclude inference of the direction or magnitude of changes in bacterial abundance ^37^, even though this is of primary interest to researchers exploring the microbiome and its impact on host fitness ^25,38,39^. Compositional methods are thus poorly suited to studies of the population biology of microbiome-dwelling bacterial taxa when bacterial load varies or when microbial fitness is a desired response variable.

Direct bacterial quantification ^25^, as well as controlled DNA spike-ins ^40^, can correct for biases and ambiguities inherent to compositional analyses. Our study provides an additional and novel framework for enabling standard high-throughput 16S sequencing approaches to provide quantitative measures of bacterial abundance when canonical approaches (e.g., qPCR) are infeasible due to host organelle contamination. Rather than being discarded, 16S read counts derived from host organelles—once curated—can provide an internal reference population against which the proportionality of other taxa can be measured ^41^. While we have established the usefulness of our estimator of bacterial load (*γ*) using a paired culture-based experiment, this need not be the only way. Bacterial culturing is an intrinsically noisy means of enumerating bacteria, due to dilution and/or counting noise. In addition, chemically-mediated antagonism or facilitation among bacterial species can cause over- or under-detection of particular combinations of taxa on agar plates ^42^. These limitations will no doubt set a lower limit to the resolution of biological effects one is capable of detecting with culture-dependent methods. Testing the generality of our approach across other plant–microbe systems, and with other means of enumerating bacteria in samples, is therefore a priority.

### Conclusions

Our study emphasizes that large effects on the population biology of *P. syringae*, and many other lineages of leaf-colonizing bacteria, may stem from the action of insect herbivores. Biotic interactions such as herbivory are absent from the classic ‘disease triangle’ of plant pathology. The role of insect herbivores in *P. syringae* epidemiology—and plant–microbiome relations in general—has been under-appreciated. Variation in bacterial abundance across samples, and the implications of relative abundance changes for bacterial fitness, are not easily detectable via compositional analyses applied to 16S data, which typically do not utilize external or internal standards. Thus, studies aiming to decipher why plant microbiomes differ in structure or function should endeavor to quantify bacterial loads in order to retain this important axis of variation as a focal response variable, while also considering additional biotic interactions commonly encountered by the hosts under study.

## Acknowledgements

PTH and NKW gratefully acknowledge funding from the National Science Foundation (DEB-1309493 to PTH, DEB-1256758 to NKW), the National Institute of General Medical Sciences of the National Institutes of Health (R35GM119816 to NKW), as well as the Rocky Mountain Biological Laboratory. We are indebted to field assistance provided by Heather Briggs, Kara Cromwell, Aaron Koning, Lucy Anderson, Kyle Niezgoda, Devon Picklum, and Nicolas Alexandre; bioinformatics advice from Tim K O’Connor; and laboratory assistance from Hoon Pyon and Amir Abidov. We thank our contacts at Argonne National Laboratory Sarah Owens and Jason Koval for their technical expertise and support.

---

**Box 1: Devising an estimator of bacterial load from 16S data**.

### Defining the estimator

To establish an estimator of bacterial load using 16S sequence data, we hypothesized that the composition of the sequencing data, in terms of host-versus bacteria-derived 16S reads, may provide information about the underlying density of bacteria. This occurs, we reasoned, because DNA templates of the two sources compete as targets during the amplification reaction, and biases towards one or the other will accrue exponentially. By this logic, the logarithm of the relative abundance of bacteria–to–host 16S counts captures information about the density of bacterial cells in the starting material. Accordingly, for each sample, we calculated the following estimator

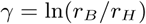

where *r*_*B*_ and *r*_*H*_ are the read counts of bacteria- and curated host-derived 16S counts for a given sample, respectively. *r*_*B*_ can be calculated at any taxonomic level, ranging from the single bASV to all of the bacteria present in the sample, by summing the sequence counts at the desired taxonomic scale.

### Validating and deploying the estimator

**Step 1: Collect paired tissue samples and count bacteria independently**.

We validated this estimator empirically by examining the relationship between *γ* and an independent measure of bacterial abundance in leaf tissues derived from bacterial culturing of a subset of the samples from the EL study. These samples were surface-sterilized, homogenized, and plated on non-selective King’s B media to enumerate bacterial colony forming units (CFU) per g starting leaf material, following Humphrey et al. [2014]. This approach is appropriate because a majority of bacterial taxa typically found to colonize leaf tissues can be cultivated in the laboratory on rich media [Lebeis et al., 2015; Humphrey et al., 2014].

**Step 2: Quantify relationship between 16S data and bacterial load**.

We then estimated the slope and intercept of the relationship between observed log_10_ CFU g^−1^ leaf tissue (hereafter log CFU) and the predictor variable *γ* for this sample set using a Bayesian linear regression, which allowed us to directly incorporate uncertainty in model fit into downstream analyses. We found a clear positive association between *γ* and log CFU, validating our usage of *γ* as an estimator of absolute bacterial abundance in leaf tissues.

**Step 3: Model relationship between *γ* and herbivore damage**.

We then deployed the validated estimator to test whether bacterial abundance as measured by *γ* was elevated in insect-damaged plant tissues. To begin, we modeled how *γ* varied across herbivore-damaged and undamaged leaves for various bacterial taxa. The illustrating example on the right shows that the distributions of *γ* calculated for all bacteria are elevated in herbivore-damaged bittercress tissues sampled from both sites EL and NP.

**Step 4: Transform results for *γ* into predicted bacterial load**.

Finally, we used posterior draws of parameters from the Step 2 model to predict how variation in *γ* translates into predicted bacterial load as expressed in log CFU. To the right, we can see that elevated *γ* in herbivore-damaged tissues translates into higher bacterial loads when predicted based on the relationship between *γ* and log CFU. Further details on how we specified and estimated models, as well as how we incorporated parameter uncertainty throughout this approach, can be found in *Methods: Quantifying & modeling bacterial abundance patterns*.

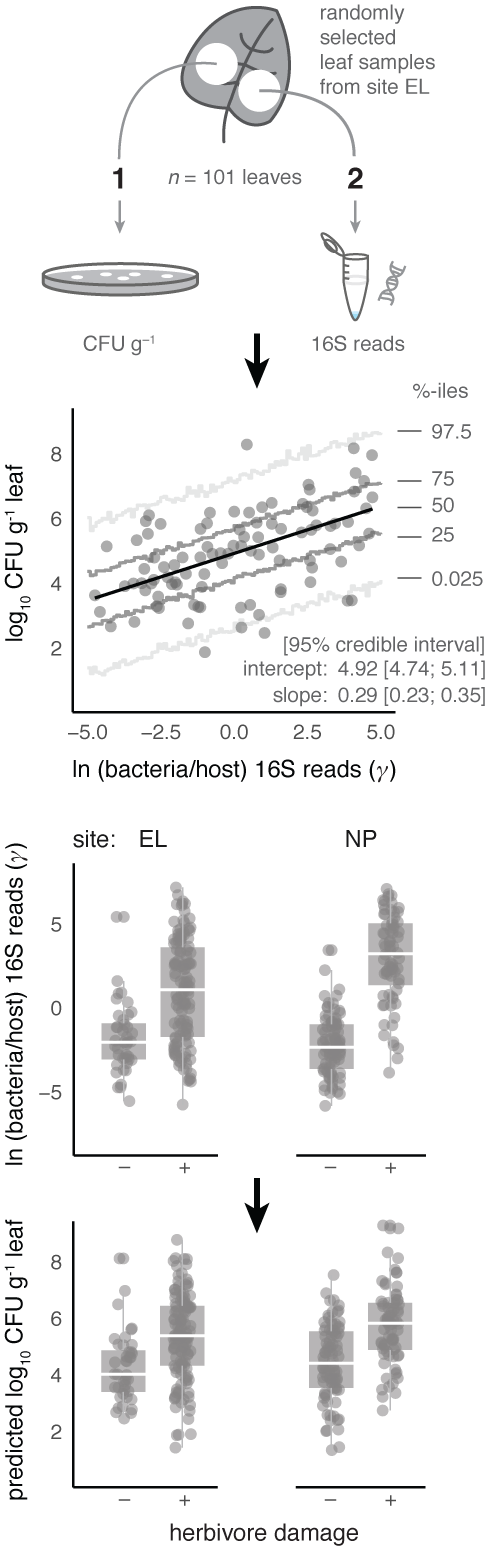

