## Supplemental online material for "Insect herbivory reshapes a native leaf microbiome"

***See*** `https://github.com/phumph/coinfection` ***for all scripts and data.***

### Supplemental Methods

#### A. Field studies.

***A.1. Hormone treatments in the field.*** Equal numbers of bittercress plots were randomized to receive either mock (control), JA, or SA treatment. One half-plot (patch; figure S1e) within each plot was subsequently randomized to receive the treatment based on the outcome of a single Bernoulli$(0.5)$ trial. We delivered hormone treatments to bittercress patches by spraying 50 mL per patch (1 mM solution of either JA or SA in 0.045% v/v methanol; Sigma–Aldrich) during mid-season prior to substantial herbivory at site EL in 2012 and at site NP in 2013, as previously described (1). For site NP, we treated plots between July 12–15, 2013, and we revisited all plots for herbivory and vegetative trait surveys as well as tissue collections between August 13–18, 2013.

At EL, we collected a single leaf from a randomly determined leaf position along each of four stems randomly selected from the 16 stems surveyed per patch. We excluded the top two leaves from sampling, which were generally $< 1\text{cm}^2$—too small to offer adequate amounts of leaf material for DNA extraction. Leaves along each stem were included irrespective of herbivore damage status. Thus, leaves with herbivore damage in our leaf sample set from EL reflected the leaf-level prevalence of herbivore damage in the EL bittercress population.

At site NP, we also measured 16 stems per patch, where four sets of four stems were identified as being closest to the inner corners of a $2 \times 2$ grid (0.5 m per side) placed in the center of each bittercress patch (figure S1e). We sampled a damaged an undamaged leaf from one of the four stems in each grid square (via randomization) and pooled the excised leaf discs from each into damaged and undamaged tissue pools. Thus, each patch contributed two samples: one containing four leaf discs from undamaged leaves and one containing four leaf discs from damaged leaves. Leaves were snipped from plant stems and held at 4 C for up to 12 h prior to processing in the laboratory.

***A.2. Sample preparation for bacterial detection.*** Leaf samples were weighed, surface-sterilized, combined into 1.5 mL microcentrifuge tubes, and flash frozen in liquid nitrogen vapor for subsequent homogenization and DNA extraction. Surface sterilization entailed a 5 s rinse in 95% ethanol, 30 s in 70% ethanol, 30 s in 10% bleach, followed by three 2 minute washes in sterile water. Tissue samples were briefly air-dried on sterilized bench-top (5 min) before being homogenized. Sterilized leaf discs were then homogenized in 2.0 mL microcentrifuge tubes in $\sim 350\mu$L 10 mM $MgSO_4$ using a TissueLyser (QIAGEN) run at max speed (50 Hz) for up to 60 s. We placed a sterilized 5 mm stainless steal ball within each tube to facilitate tissue disruption.

We then extracted DNA from all leaf homogenate using the ThermoFisher PureLink Genomic DNA extraction kit following standard protocols. We eluted DNA in 35 $\mu$L of sterile water and shipped 10 $\mu$L aliquots to Argonne National Laboratory (ANL) at 4 C. ANL staff then performed PCR amplification of the V4 region of 16S using published primers (4, 5) followed by Illumina MiSeq sequencing according to Earth Microbiome Project (EMP) protocols (2) (see `http://press.igsb.anl.gov/earthmicrobiome/protocols-and-standards/16s/` for full details). To reduce amplification of chloroplast- and mitochondrial-derived 16S, we amended the amplification reaction mixtures with two peptic nucleic acid (PNA) oligomer PCR clamps at a final concentration of 1.25 $\mu$M each (3).

#### B. Analysis of bacterial abundance patterns.

***B.1. Overview.*** Our statistical objective was to assess the evidence that herbivore damage is associated with systematic differences in the abundance of endophytic bacteria compared to undamaged leaves. Below we describe how we modeled the abundance of bacteria by assessing the proportionality between bacterial reads and host-derived reads across the sample set ($\gamma$; see **Box 1** in main text for details). We supplied all non-zero entries for each bASV to the models below, discarding any ASV seen in only one library or with fewer than 10 counts across the dataset. At no point did we perform down-sampling (i.e., rarefaction) on 16S libraries.

***B.2. Curating host-derived 16S reads.*** To curate host-derived reads, we first assigned provisional taxonomy to all ASVs in the datasets via the Ribosomal Database Project Bayesian classifier (6) to closely inspect putatively plant- or fungus-derived sequences. All such ASVs matching non-Brassicaceae plant- or fungi-derived 16S were discarded, while those annotated as *Cardamine* (for chloroplast) or *Arabidopsis* (for mitochondria) were retained. We then manually verified monophyly of putative bittercress-derived ASVs after constructing maximum likelihood trees of all non-bacterial ASVs, using **R** package `ape` (7). We then pooled bittercress-derived chloroplast and mitochondria reads as 'host-derived' for downstream calculations of bacterial abundance (below).

***B.3. Details of Statistical Models.*** *Stage 1: $\gamma$ models.* As the leaf sample-level response variable, 16S counts were expressed relative to host-derived counts, and this ratio was log-transformed to derive a proportionality measure for each bASV: $\gamma = \ln(r_B/r_H)$, where $r_B$ and $r_H$ are the read counts of bacteria- and curated host-derived 16S counts for a given sample, respectively (see **Box 1**, main text for details). Our choice of base $e$ in the logarithms was made by convention, and alternate logarithm bases will not impact how $\gamma$ behaves mathematically.

For each bacterial family, we ran a series of increasingly flexible hierarchical Bayesian (log) normal regression models with observed $\gamma$ as the response variable for data collected at sites EL and NP separately (model definitions given below). We included group-level model terms to capture variation in the intercept and slope of the relationship between $\gamma$ and herbivore damage, and we also included terms to model heterogeneity of variances between sample classes as well as overall skew. We compared models using approximate leave-one-out Bayesian information criterion (LOO-IC) (8), and we heuristically took the model displaying the lowest LOO-IC as the candidate best-fitting model for downstream purposes.

From each candidate best stage-1 model, we generated posterior predicted values of the response variable ($\tilde{\gamma}$) by conducting 200 replicate simulations, each the size of the original input data, from the posterior distributions of model parameters. This posterior predictive distribution of the input data $p(\tilde{\gamma}|\gamma,\theta)$ represents the range of values that the response variable is likely to take under the specified model, and it reflects both uncertainty in the parameter estimates ($p(\theta|\gamma)$) from the model as well as the extent of group-level and residual variation present in the data itself.

*Stage 2: absolute abundance models.* Vectors of $\tilde{\gamma}$ were used as the input predictor variable in a model describing the relationship between $\gamma$ and bacterial abundance on the scale of logCFU. When combined with estimated regression parameters (also drawn from their posterior distributions, for each simulated leaf sample separately), this transforms input $\tilde{\gamma}$ values into estimates of absolute bacterial load. Regression parameters for the abundance model were estimated from a sample set from which bacteria were enumerated by culturing as well as 16S sequencing; full details on establishing the relationship between $\gamma$ and absolute bacterial load are given in **Box 1** of the main text.

We used these posterior predicted estimates of logCFU to calculate the difference in abundance of each bacterial family between damaged and undamaged leaf sets. To calculate abundance values at the higher taxonomic scale of family, we summed the predicted counts (on the linear scale) of the individual bASVs at the level of the leaf sample for each of the 200 posterior simulations separately. Including a bASV in the family-level sums with an observed count of zero for a particular leaf sample was determined probabilistically after modeling whether the predicted bASV abundance would permit detection given the detection threshold for each particular sample (see section ***Accounting for sample-level detection thresholds***, below).

We then computed the difference in median abundances between damaged and undamaged leaf sets. We did not use means of predicted log CFU values because of the potentially large influence of positive outliers on the log scale on summary statistics calculated on the exponentiated scale (i.e., $\text{med}(\log x) = \log \text{med}(x)$, while $\log \text{E}[x] \neq \text{E}[\log x]$). We expressed the median difference in predicted bacterial density between leaf types in units of $\log_2$ so that we could represent effects in terms of the number of bacterial doublings (i.e., cell divisions) that comprise the predicted abundance differences between sample types. This calculation was repeated for each of the 200 replicate posterior draws to generate the posterior predictive distribution of the difference in within-host density for bacteria in damaged versus undamaged leaves.

***B.4. Model definitions.*** Stage 1 models of $\gamma$ estimate the population-level effect of herbivore damage, as well as among-bASV-level variation in the intercept (i.e., variation in abundance among un-damaged leaves) in addition to the slope of the difference between un-damaged and damaged leaves. This model structure imposes the prior assumption that bASV abundance patterns are drawn from Gaussian distribution around a family level mean intercept and/or slope values and is represented below as the 'base' model (**ga3**):

$$\begin{aligned} \gamma|\alpha,\beta,\sigma &\sim \mathcal{N}(\alpha_j + \beta_j D, \sigma_0) \\ \alpha_j &\sim \mathcal{N}(\alpha_0, \tau_\alpha) \\ \beta_j &\sim \mathcal{N}(\beta_0, \tau_\beta) \end{aligned} \tag{ga3}$$

In the model above, $\alpha_j$ represents the intercept term for bASV $j$, which is centered around the overall mean intercept $\alpha_0$ and deviates from it by an amount given by the standard deviation term $\tau_\alpha$. Similarly, $\beta_j$ represents the *difference* in abundance of a given bASV $j$ in damaged leaves, which is centered around a family-mean slope $\beta_0$ and deviates from it by an amount given by the standard deviation term $\tau_\beta$. $D \in \{0,1\}$ and reflects the damage status of a given leaf. Finally, the standard deviation term $\sigma_0$ controls the magnitude of residual variation around each bASV-level estimate of abundance in both damage classes. Priors on population-level terms, $p(\alpha_0, \beta_0, \sigma_0)$, and group-level hyperpriors $p(\tau_\alpha, \tau_\beta)$ were set to be weakly informative, and we thus retained default prior distribution functions in `brms` to achieve this (9, 10).

We also included additional model parameters to empirically improve fit. To do this, we introduced an additional residual error term to model **ga3** that accounts for heterogeneity of variances between leaf damage classes, yielding model **ga4**:

$$\gamma|\alpha,\beta,\sigma \sim \begin{cases} \mathcal{N}(\alpha_j + \beta_j D, \sigma_1), & \text{if D=1} \\ \mathcal{N}(\alpha_j + \beta_j D, \sigma_0), & \text{otherwise} \end{cases} \tag{ga4}$$

We then simplified models **ga3** by removing the group-level slope term ($\beta_j$) and simply estimated a single family-level $\beta_0$, yielding model **ga1**, which was further simplified by removing the slope terms entirely (i.e., intercept-only model, **ga0**). Model **ga2** is identical to model **ga1** but includes the additional residual $\sigma$ term per damage class, as in the definition of model **ga4** above.

Finally, a direct analog of each of these models was estimated using instead skew-Normal sampling distribution for the likelihood, where a hyperprior on the skew-Normal shape parameter $\delta$ was given as $p(\delta) \sim$ half-$\mathcal{N}(0, 0.5)$ (by default).

We fit all model with Stan with its interface to **R** (11) via `Rstan` (12) using package `brms` (9, 10). We estimated each model described above using four chains of 8000 iterations each with a burn-in fraction of 0.5. We monitored Markov Chain mixing properties via standard measures implemented by default in `brms` and then compared models via calculating approximate leave-one-out (LOO) information criterion (8) implemented in `brms`. For each family, the model showing the lowest LOO-IC was further inspected via posterior predictive model checks prior to model selection for reporting.

***B.5. Posterior predictions of bASV abundance.*** To generate predicted bASV absolute abundances, we drew coefficients from the joint posterior distribution of parameters for the log-ratio model and used them to estimate the posterior predicted values of the response variable, $\tilde{\gamma}$. The simulated $\tilde{\gamma}$ values were then fed as predictors into the abundance-model:

$$\log_{10}\widetilde{\text{CFU}}\ g^{-1}|\alpha,\beta,\sigma \sim \mathcal{N}(\alpha + \beta\tilde{\gamma}, \sigma^2) \tag{abundance model}$$

Parameters for the abundance model were re-drawn for each posterior predictive data point such that parameter uncertainty was incorporated at both modeling stages. This procedure was repeated 200 times for simulated datasets of the same size as the actual sample. We summed bASV-level abundances (on the linear scale) that were predicted by these simulations at the taxonomic level of bacterial family after correcting for detection biases (see below).

***B.6. Accounting for sample-level detection thresholds.*** Taking sums across bASVs within families required us to correct for uncertainty in the presence/absence of particular bASVs, as many bASVs had observed counts of zero across several leaf samples. To decide whether to treat each zero count as a true or false absence, we estimated whether the sample-wide predicted abundance of a given bASV across leaves would lie above or below the empirical detection threshold for each individual leaf sample. The detection threshold for each leaf is set by the total count of host-derived 16S—not by the total library size. We calculated such sample-level thresholds for each leaf as $\gamma_T = 1/r_H$. Ultimately this threshold is set by sequencing depth, but only indirectly: instead, it is the number of host-derived reads per sample that sets the ability to detect a bASVs at a given level of $\gamma$. Even for samples with many sequencing reads, high-abundance bASVs can cast statistical shadows on low-abundance bASVs by driving down the count of host-derived 16S. Thus, samples for which certain bASVs are highly abundant make it more difficult to detect bASVs with lower $\gamma$ in that particular sample.

To make such prevalence corrections, we calculated the sample-wide distribution of $\tilde{\gamma}$ for each individual bASV across each leaf sample type (damaged and undamaged) using 1000 posterior draws from the candidate best stage-1 model for each bacterial family. We then calculated $\gamma_T$ for each sample that had an observed zero count for the focal bASV. We scored each observed zero count as 'true' (i.e., truly absent) if the detection threshold for each sample, $\gamma_T$, was below the lower 1%-ile of the posterior predicted abundance distribution $p(\tilde{\gamma}|\gamma)$. Heuristically, a zero counts for a given sample was scored as 'false' (i.e., a missed detection) when $\gamma_T$ fell anywhere within $p(\tilde{\gamma}|\gamma)$ above the lower 1%-ile. This approach avoids including high-abundance bASVs from samples when there is evidence against its presence, while also avoiding the neglect of low-abundance bASVs whose abundance likely lies at or below detection limits given our sampling depth. Implicit in our approach is the assumption that observed zero counts of on-average low-abundance bASVs are more likely to be missed detections rather than true absences.

***B.7. Applying a finite sample size correction.*** To further reduce our potential bias towards over- or under- inclusion of bASVs in family-level totals, we addressed finite sample size bias by estimating sample-wide bASV prevalence. The main effects of this procedure are to relax the assumption that a given bASV found in 100% of our leaf samples has an actual population-wide prevalence of 100%, as well as to admit that rarely-detected bASVs may in fact have a prevalence greater than that observed. We implemented these notions by modeling bASV prevalence using the posterior Beta distribution with the weakly-informative Jeffrey's conjugate prior. This posterior Beta distribution has the parameters $\alpha = 0.5 + k$ and $\beta = 0.5 + n - k$, such that uncertainty in prior prevalence expressed through Jeffrey's prior ($\alpha = \beta = 0.5$) gives posterior prevalence estimates always less than 1.0. bASVs with a sample prevalence of zero were already excluded from this analysis by virtue of their never having been detected in the first place. We generated prevalence estimates by drawing from the posterior Beta distribution for each bASV. A Bernoulli random variable was then drawn based on this value in order to determine whether to include the predicted bASV abundance in a given simulated family-level summation for a given leaf sample. Family-level totals are thus probabilistic sums of bASVs weighted by their predicted prevalence, which incorporates uncertainty in presence/absence arising form finite sampling at the level of individual sequencing libraries as well as the number of total libraries sequenced.

**C. Estimating compositional changes in leaf bacterial communities.** We examined how differential growth under herbivory of distinct members of the bacterial community would impact overall signatures of ecological diversity and similarity. Using our predictive posterior distributions of bacterial abundances (i.e., median log CFU), we calculated standard ecological summary statistics of community-level compositional variation across herbivore-impacted versus non-impacted leaves (Shannon evenness $J'$, and Shannon–Jensen divergence $SJ$). We used the vector of 200 simulated median predicted abundances (on the linear scale) for damaged and undamaged leaf sets and transformed these values into relative abundances. We then calculated the posterior predictive distribution of $J'$ using the 200 simulated damaged and undamaged communities, compared the distribution of differences between leaf classes, and finally examined evidence for community-level differentiation via $SJ$ divergence. We repeated this procedure for the 200 simulated *Pseudomonas* spp. communities arising from growth rate data derived from our greenhouse experiment (described below). Evenness was calculated as:

$$J' = \Big(-\sum_{i=1}^{S} p_i \log p_i\Big) / \Big(\log S\Big)$$

where $p_i$ is the individual relative abundance of a given taxonomic group, and $S$ is the total number of such groups in the sample.

Shannon–Jensen divergence was calculated as the average Kullback–Leibler divergence (i.e., $D(a||b)$) between each of the component relative frequency vectors and the vector comprising their average:

$$SJ(X||Y) = \frac{1}{2}\Big(D(X||M) + D(Y||M)\Big)$$

where $M = 0.5 \times (X + Y)$ and vectors $X$ and $Y$ contain relative frequencies of taxa present in each of the two focal samples.

**D. Modeling within-host performance of *Pseudomonas* spp. in experimental infections.** Using a Bayesian regression model, we estimated the effect of hormone pre-treatment (Mock versus JA) on bacterial doublings (i.e., cell divisions) after 2 d of growth within bittercress leaves following syringe inoculation in the greenhouse, as in (1). We estimated separate intercept and slope terms for each of the twelve *Pseudomonas* spp. strains used in this experiment with a common residual variance, as well as a plant ID grouping term to account for intercept-level variation across replicate plants ($n = 32$ separate plants total). Further details of experimental design can be found in the main text.

**E. Modeling variation in herbivore damage in the field.** We implemented Bayesian Gamma–Poisson (i.e., negative binomial, NB) mixture models to estimate how damage levels from *S. nigrita* leaf miners varied across bittercress patches at site NP in relation to plant defense hormone pre-treatments. We summed the counts of mined leaves per per patch ($n = 16$ stems per patch) and took this as the response variable, which was modeled as a function of patch-level treatment (Mock, JA, or SA). We included a plot-level grouping term (i.e., random effect), which imposes that variation in the average (log) response variable per plots is normally distributed. We also included an offset term to account for different sampling depth (i.e., [log] number of leaves summed across the $n = 16$ stems per patch). Hyperpriors on the mixture component for NB models were set as `brms` defaults.

**F. Modeling fruit set across bittercress patches in the field.** We used a Bayesian NB model to estimate how patch-level fruit set relates to the intensity of patch-level *S. nigrita* leaf miner damage under field conditions at site NP. We took the summed number of fruits per patch as the response variable, which was modeled as a function of the total number of leaves, the total number of leaves with leaf miner damage, and the average stem height across the $n = 16$ stems sampled per patch. We also included plot-level grouping term, which implicitly models among-patch variation in (log) baseline fruit set as normally distributed. We report the model without the total number of leaves term, as this term showed a posterior distribution symmetric around 0, and dropping it from the model was supported by LOO-IC.

#### Supplemental Figures and Tables

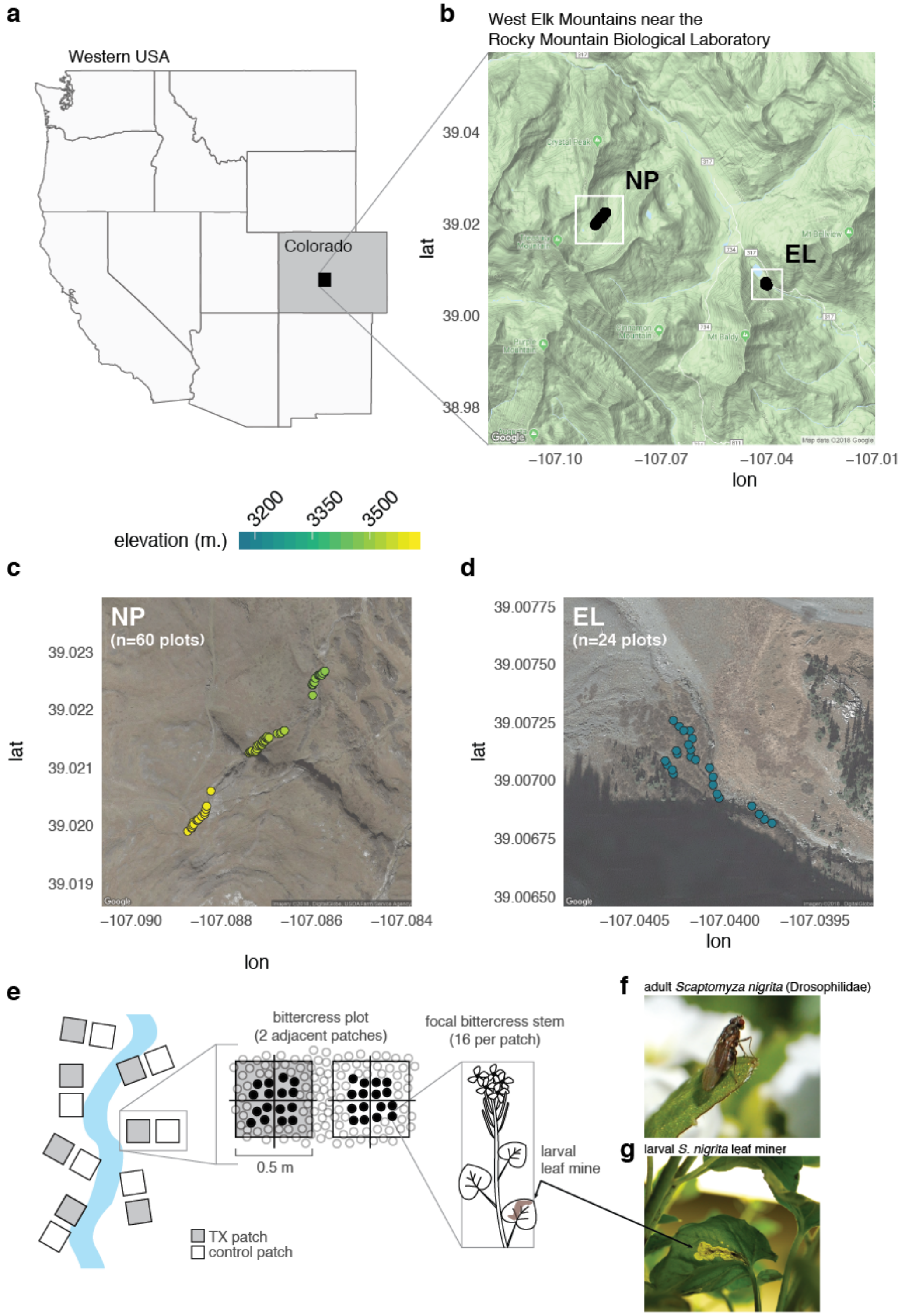


**Fig. S1. Location and design of field studies on plant co-infection by herbivores and bacteria**. **a–b.** Regional map showing location of field sites near the Rocky Mountain Biological Laboratory in the West Elk Mountains of Colorado, USA. **c–d**. Field transects in bittercress populations at sites Emerald Lake (EL, 2012) and North Pole Basin (NP, 2013). **e**. Schematic of transect, plot, and patch-level sampling design. Focal bittercress stems were surveyed for total leaves, stem height, number of mined leaves, and number of fruits (NP only) following early season exposure to exogenous plant defense hormones jasmonic acid (JA), salicylic acid (SA), or a mock control solution (see Methods). **f**. *Scaptomyza nigrita* adult female fly, which lays eggs within punctures created in bittercress leaves using their dentate ovipositors. **g**. Larval *S. nigrita* create mines within bittercress leaves after hatching and complete development within leaf tissues, typically on a single host plant.

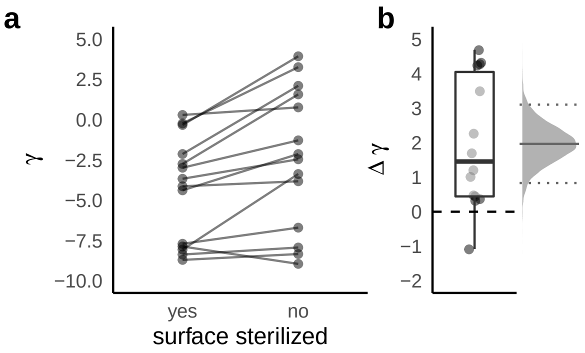


**Fig. S2. Surface-sterilizing leaves prior to DNA extraction reduces bacterial abundance**. **a.** Paired leaf discs from $n = 18$ samples from an independent sample set from the EL study site shows consistent reduction in a metric of bacterial abundance ($\gamma$) calculated from 16S sequence data (see Box 1, main text). **b.** Distribution of observed differences in $\gamma$ between non-sterilized and sterilized leaf discs, with the adjacent density plot representing the posterior probability of the difference from a Bayesian regression model. The density plot is annotated with the median (solid line) as well as the 2.5%, and 97.5% quantiles of the posterior distribution of the estimated difference.

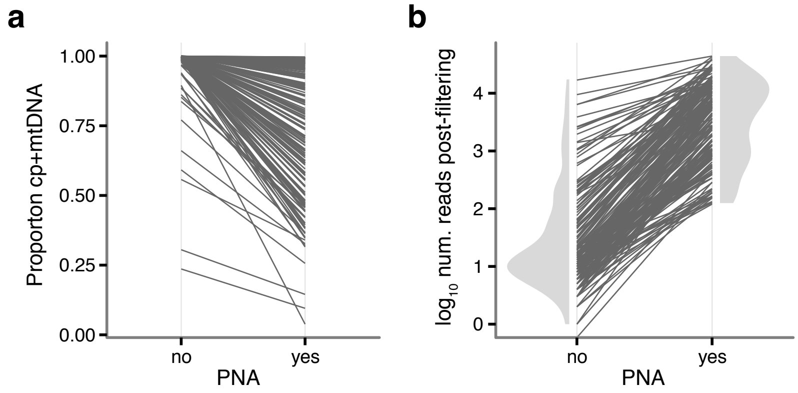


**Fig. S3. Host-derived 16S reads dominate sequencing libraries without addition of peptic nucleic acid (PNA) PCR clamps. a.** Proportion of 16S reads flagged as host-derived (chloroplast, cp and mitochondrial, mt) with and without inclusion of 1.25 $\mu$M PNAs in 16S amplification reactions. **b.** Total ($\log_{10}$) reads of bacteria-derived 16S counts with and without addition of PNAs ($n = 384$ libraries)

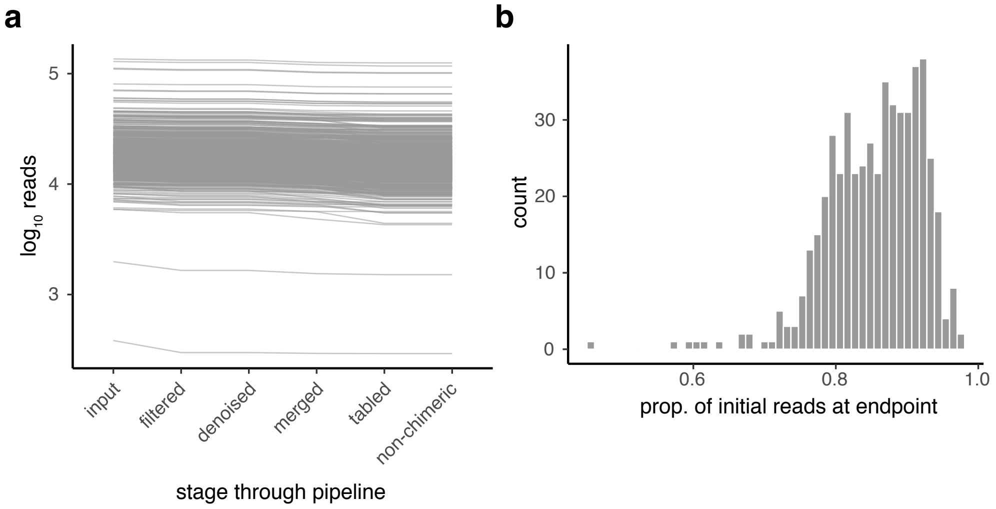


**Fig. S4. Summary of sample processing and quality control pipeline of `DADA2`. a.** Counts of 16S reads per sequencing library passing each step of the pipeline. **b.** Proportion of input reads surviving post-processing per library.

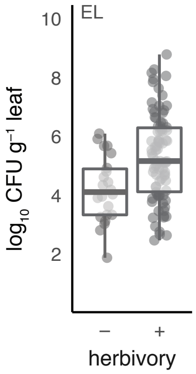


**Fig. S5. Culturable bacteria loads are higher in herbivore-damage leaves within the calibration sample set from site EL** ($n = 101$; Welch's unequal variance $t$-test $t = 3.86$, $p < 0.001$). Magnitude of increase in damaged leaves for culturable bacterial loads ($3.7[1.8 - 5.6$ 95% c.i.$]$ bacterial doublings) closely mirror those found for all Bacteria using 16S sequence data at site EL (table S2) as well as the findings reported for a parallel and independent transect survey of bittercress leaves from site EL published by Humphrey et al. (1).

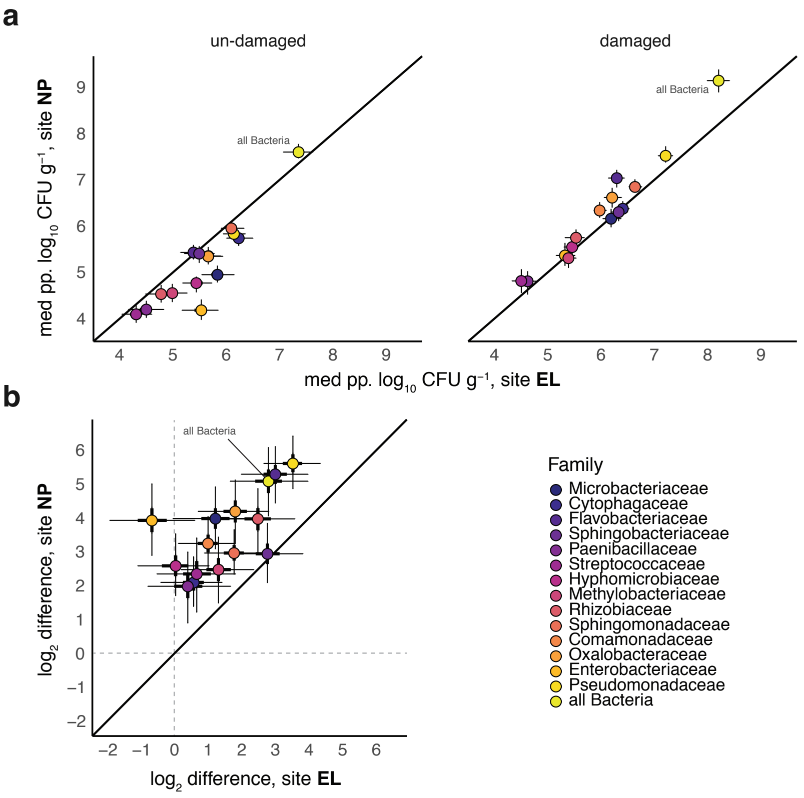


**Fig. S6. Correspondence between predicted bacterial abundance and herbivory effects between sites EL and NP**. **a.** Plotted are median (circles) $\pm 95\%$ posterior distributions of predicted abundance for bacterial bASVs summed at the family level for sites EL ($x$-axis) and NP ($y$-axis). **b.** Comparison between the magnitudes of $\log_2$-fold differences between damaged and undamaged leaves at sites EL ($x$-axis) and NP ($y$-axis). Middle 50%-ile and 95%-iles of median effects (circles) are depicted by thick and thin bars, respectively. On both plots, we also show data summed across all taxa in the dataset ('all Bacteria').

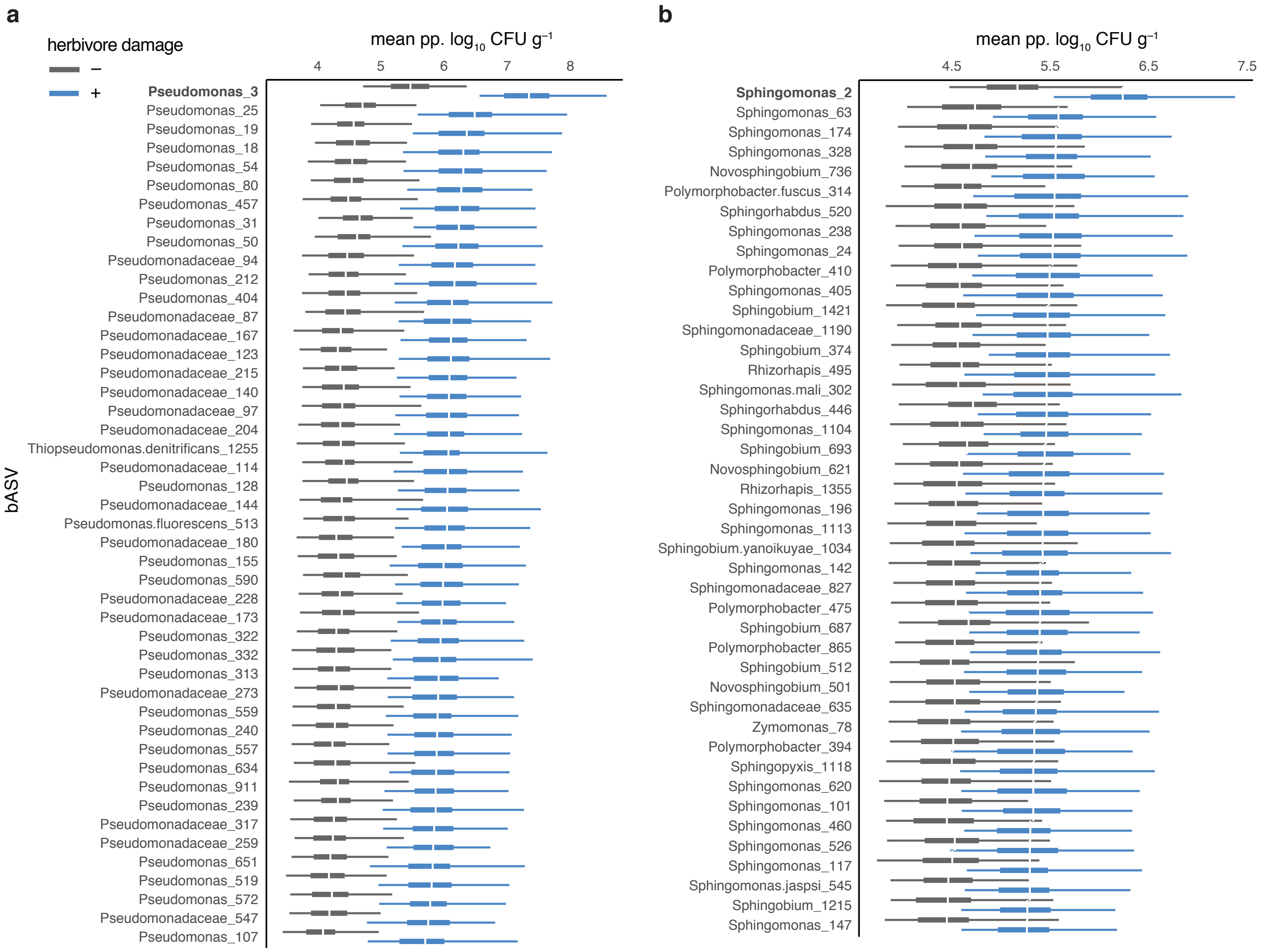


**Fig. S7. Variation among bASVs in slope and intercept parameters across damaged and un-damaged leaves from site NP**. Data are posterior predicted bacterial abundance from two-stage modeling procedure (see Methods) plotted at the bASV level for the families Pseudomonadaceae (**a**) and Sphingomonadaceae (**b**). Results derive from models which included random slope terms for bASV-level variation in effect of herbivory (term $\tau_\beta$ of models *skn3* and *skn4*, table S2). Bolded bASV names at the top represent those within each group with substantially elevated intercept and slope estimates.

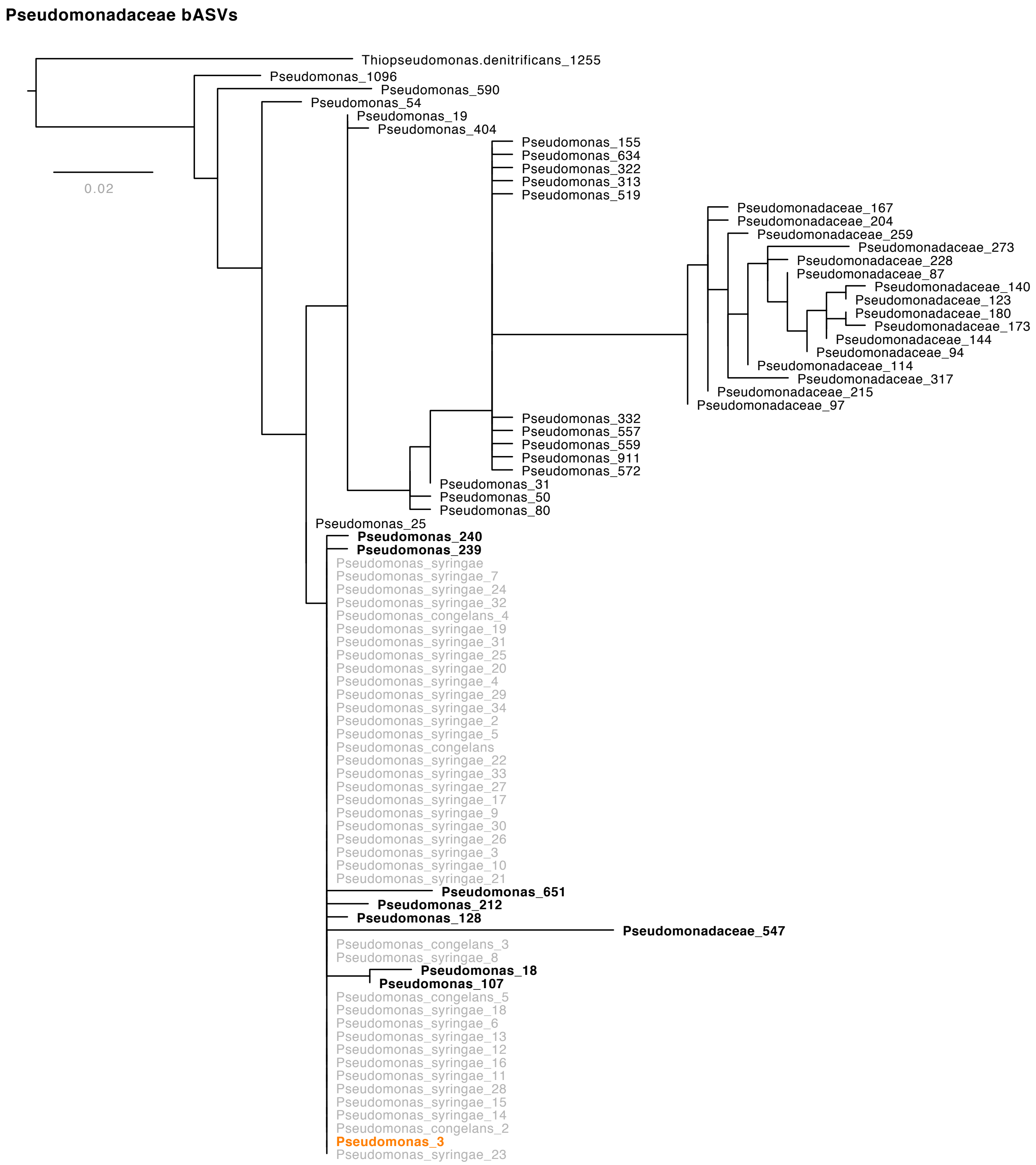


**Fig. S8. Phylogeny of aligned 16S sequence with various *P. syringae* accessions showing abundant bASVs nesting within this clade**. The most abundant bASV across our sample set, `Pseudomonas3`, clades with *P. syrinage*. `Pseudomonas3` was mapped to the putatively phytopathogenic *P. syringae* species complex via maximum likelihood estimation of a phylogenetic tree using RAxML 8.2, (13) using all *Pseudomonas* bASVs ($n = 49$) and their top ten non-redundant BLAST hits.

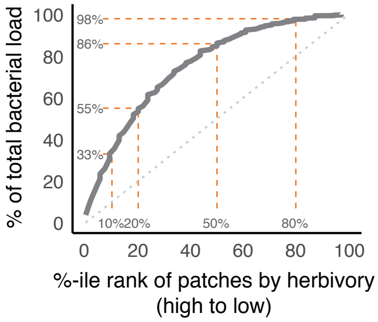


**Fig. S9. Plant patches with high herbivory harbor a disproportionate fraction of the estimated population of the most abundant *P. syringae* bASV**. Scaling of percentile rank (high to low) of patch-level herbivore load with total population-level percentage of bacterial propagules present in the sampled patch. At site NP, the top 20% of plant patches with the most herbivory harbor $>$50% of bacterial propagules in the plant population.

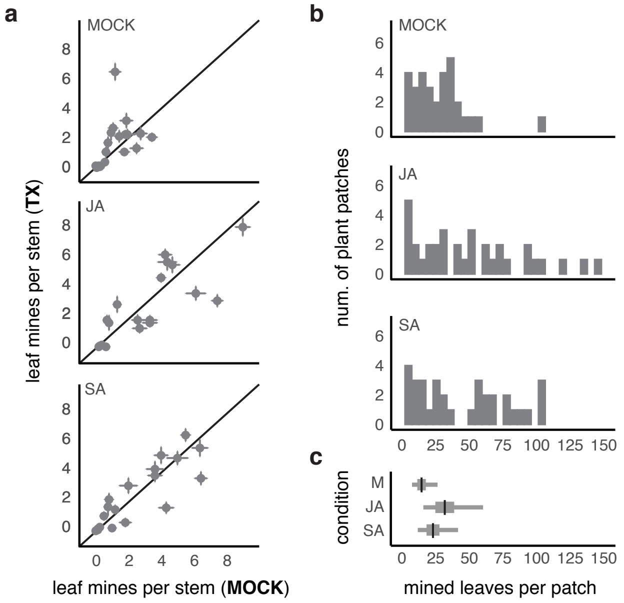


**Fig. S10. Observed distributions (a–b) and estimated marginal effects (c) of hormone pre-treatment effects on levels of *S. nigrita* herbivory in bittercress populations at site NP**. **a.** For each plot separately, plotted is the average ($\pm 1 \times$ std. error) leaf mines per stem calculated at the patch level ($n = 16$ stems per patch) for mock-treated ($x$-axis) versus hormone-treated ($y$-axis) patches. The three panels represent plots assigned to each of the three conditions: mock (i.e., control), jasmonic acid (JA) or salicylic acid (SA). **b.** Histograms of patch-level leaf miner damage broken down by patch-level treatment. **c.** Marginal effects for estimates of patch-level treatment on total mined leaves per patch (see table S5 for statistical results). Black bars are posterior means, while thick and thin bars comprises middle 50%- and 95%-iles of posterior distributions of model terms marginalized over all other parameters.

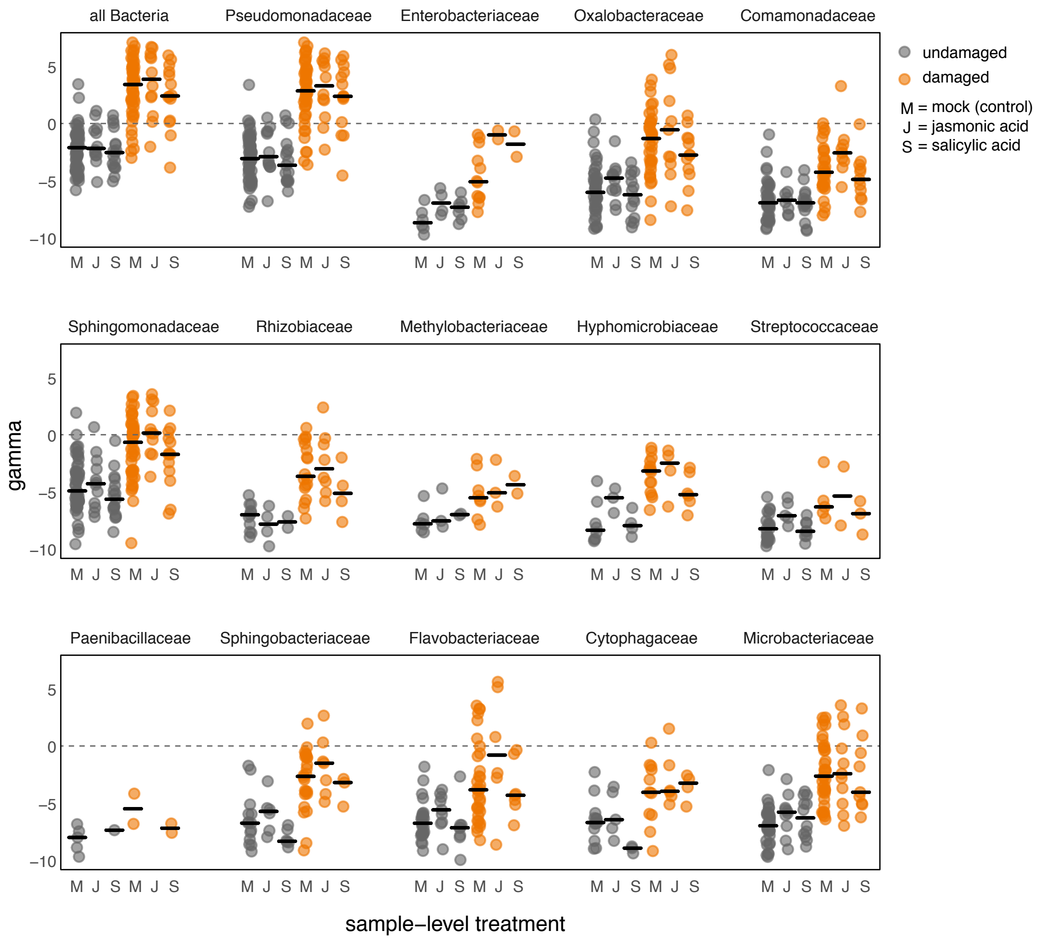


**Fig. S11. Hormone pre-treatment (Mock, JA, or SA) five weeks prior to sampling does not leave a clear signature in the distribution of $\gamma$ across samples for undamaged (gray) or herbivore-damaged (orange) leaf samples**. For each of the 14 most abundant bacterial families, in addition to all Bacteria, we have plotted the distributions of raw $\gamma$ for damaged and undamaged samples across mock- (M), JA-, or SA-treated plant patches. Black bars represent medians for the respective distribution, and data points are slightly jittered along the $x$-axis. Systematic differences in $\gamma$ can be easily seen between damaged versus undamaged sample classes, whereas no systematic differences can be seen between the different hormone treatment classes within each damage class. If any effects of hormone treatments are indeed present, they do not constitute a discernible feature of these data, supporting the choice to devote minimal attention to this aspect of our experiment.

**Table S1.** Model coefficients linking 16S counts to CFU

| Term | Estimate [95% c.i.] |
|---|---|
| $\alpha$ intercept | 4.92 [4.7;5.14] |
| $\beta$ slope | 0.29 [0.22;0.36] |
| $\sigma$ | 1.18 [1.03;1.35] |

**Table S2.** Model results for family-level bacterial abundance

| Family | model | $\alpha_0$ | $\beta_0$ | $\tau_\alpha$ | $\tau_\beta$ | $\tau_p$ | $\sigma_0$ | $\sigma_1$ | $\alpha$ |
|---|---|---|---|---|---|---|---|---|---|
| **site EL** | | | | | | | | | |
| all Bacteria | ga2 | -1.95 [-2.66;-1.26] | 2.89 [2.02;3.78] | – | – | 0.68 [0.04;1.41] | 1.94 [1.51;2.51] | 1.59 [1.17;2.08] | – |
| Pseudomonadaceae | skn2 | -7.07 [-7.58;-6.58] | 2.66 [2.2;3.08] | 0.67 [0.43;0.97] | – | – | 2.17 [1.91;2.48] | 1.41 [1.21;1.61] | 2.62 [1.92;3.47] |
| Enterobacteriaceae | ga2 | -5.91 [-8.01;-3.81] | 0.86 [-0.75;2.51] | 2.01 [0.87;3.96] | – | – | 2.6 [1.63;4.13] | 0.65 [0.34;1.09] | – |
| Oxalobacteraceae | ga1 | -6.71 [-7.96;-5.5] | 1.85 [1.03;2.66] | – | – | – | 2.95 [2.75;3.17] | – | – |
| Comamonadaceae | ga2 | -6.65 [-7.17;-6.14] | 0.67 [0.26;1.08] | 0.78 [0.44;1.21] | – | – | 1.43 [1.22;1.69] | 1.26 [1.04;1.5] | – |
| Sphingomonadaceae | skn3 | -6.95 [-7.75;-6.18] | 1.71 [0.88;2.4] | 0.97 [0.52;1.59] | 0.7 [0.18;1.4] | – | 2.52 [2.4;2.65] | – | -0.38 [-1.45;0.8] |
| Rhizobiaceae | skn3 | -7.62 [-9.97;-5.2] | 2.62 [-0.26;5.33] | 1.27 [0.04;3.96] | 2.09 [0.16;5.3] | – | 2.28 [1.8;2.93] | – | -3.25 [-8.22;0.74] |
| Methylobacteriaceae | ga2 | -6.05 [-6.97;-5.13] | 0.31 [-0.3;0.89] | 1.02 [0.49;2.02] | – | – | 1.36 [1.06;1.76] | 1.38 [1.02;1.79] | – |
| Hyphomicrobiaceae | ga0 | -5.42 [-6.22;-4.71] | – | – | – | – | 2.07 [1.81;2.38] | – | – |
| Streptococcaceae | ga2 | -7.32 [-8.08;-6.67] | 0.57 [0.08;1.08] | 0.39 [0.01;1.72] | – | – | 1.08 [0.84;1.4] | 1.37 [0.99;1.84] | – |
| Paenibacillaceae | skn1 | -6.82 [-8.03;-5.69] | 1.07 [0.02;2.19] | – | – | – | 1.57 [1.29;1.94] | – | 2.18 [-1.18;6.34] |
| Sphingobacteriaceae | skn2 | -6.68 [-7.9;-5.37] | 2.43 [1.14;3.65] | 0.9 [0.11;1.81] | – | – | 1.8 [0.99;3.16] | 1.66 [0.86;2.82] | 3.21 [1.49;5.99] |
| Flavobacteriaceae | skn2 | -7.06 [-7.96;-6] | 2.14 [1.07;3.17] | 0.67 [0.05;1.71] | – | – | 1.84 [1.29;2.64] | 1.7 [1.09;2.41] | 4.18 [1.71;7.84] |
| Cytophagaceae | ga0 | -5.2 [-5.48;-4.92] | – | – | – | – | 2.08 [1.92;2.27] | – | – |
| Microbacteriaceae | skn3 | -5.49 [-6.57;-4.5] | 1.33 [0.15;2.47] | 0.48 [0.02;1.52] | 0.8 [0.05;2.05] | – | 3.05 [2.81;3.32] | – | 2.52 [1.35;3.88] |
| **site NP** | | | | | | | | | |
| all Bacteria | skn2 | -2.2 [-2.6;-1.81] | 5.12 [4.48;5.75] | – | – | 1.28 [0.78;1.71] | 1.32 [0.94;1.74] | 1.84 [1.27;2.7] | -1.26 [-5.95;1.6] |
| Pseudomonadaceae | skn4 | -7.74 [-8.04;-7.47] | 4.79 [4.58;5] | 0.84 [0.65;1.09] | 0.33 [0.06;0.63] | – | 1.64 [1.57;1.72] | 1.65 [1.55;1.75] | 1.56 [1.21;1.92] |
| Enterobacteriaceae | ga2 | -7.74 [-8.66;-6.69] | 3.47 [2.13;4.76] | 0.69 [0.03;2.29] | – | – | 1.12 [0.79;1.63] | 2.36 [1.36;3.76] | – |
| Oxalobacteraceae | skn4 | -6.82 [-7.25;-6.46] | 3.66 [2.96;4.25] | 0.3 [0.01;0.79] | 0.56 [0.05;1.39] | – | 1.94 [1.76;2.15] | 1.47 [1.28;1.68] | 1.98 [1.03;3.06] |
| Comamonadaceae | skn2 | -7.58 [-7.97;-7.22] | 2.86 [2.38;3.34] | 0.43 [0.12;0.82] | – | – | 1.35 [1.18;1.57] | 1.62 [1.33;1.95] | 2.45 [1.39;3.79] |
| Sphingomonadaceae | skn4 | -7.25 [-7.62;-6.9] | 2.82 [2.41;3.2] | 0.67 [0.46;0.95] | 0.29 [0.03;0.65] | – | 1.81 [1.67;1.96] | 1.19 [1.04;1.34] | 1.54 [0.64;2.29] |
| Rhizobiaceae | ga2 | -7.78 [-8.88;-6.82] | 4.04 [3.31;4.75] | 1.02 [0.34;2.24] | – | – | 1.22 [0.86;1.74] | 1.63 [1.07;2.35] | – |
| Methylobacteriaceae | ga2 | -7.4 [-8.18;-6.67] | 2.22 [1.12;3.36] | 0.42 [0.01;1.39] | – | – | 1.16 [0.81;1.7] | 1.67 [0.95;2.77] | – |
| Hyphomicrobiaceae | ga1 | -7.56 [-8.33;-6.8] | 3.07 [2.21;3.93] | – | – | – | 1.61 [1.35;1.93] | – | – |
| Streptococcaceae | skn2 | -8.07 [-8.58;-7.49] | 2.07 [0.85;3.52] | 0.35 [0.01;1.49] | – | – | 1.13 [0.89;1.45] | 1.86 [1.1;3.17] | 4.95 [1.46;9.29] |
| Paenibacillaceae | ga2 | -8.19 [-9.41;-6.94] | 1.61 [-0.94;4.27] | 1.09 [0.12;2.8] | – | – | 0.67 [0.26;1.65] | 4.05 [0.89;12.7] | – |
| Sphingobacteriaceae | ga1 | -6.61 [-7.15;-6.05] | 3.44 [2.8;4.09] | – | – | – | 1.97 [1.76;2.21] | – | – |
| Flavobacteriaceae | skn2 | -7.08 [-7.42;-6.73] | 4.47 [3.87;5.1] | 0.22 [0.01;0.64] | – | – | 1.52 [1.32;1.76] | 2.01 [1.65;2.41] | 2.58 [1.34;4.15] |
| Cytophagaceae | ga1 | -6.87 [-7.49;-6.27] | 2.38 [1.66;3.09] | – | – | – | 1.84 [1.59;2.12] | – | – |
| Microbacteriaceae | skn4 | -7.31 [-7.96;-6.8] | 3.51 [2.37;4.44] | 0.53 [0.1;1.28] | 0.93 [0.09;2.35] | – | 1.43 [1.24;1.68] | 1.75 [1.43;2.12] | 1.48 [-0.52;2.91] |

*Notes:* $\alpha$ = Skew-Normal shape parameter; $\tau p$ = among-patch standard deviation (group-level term, only included in all Bacteria model).

**Table S3.** Strain-level model terms for experimental infections

| Strain | Intercept [95% c.i.] | Slope [95% c.i.] |
|---|---|---|
| ***P. syringae*** | | |
| 20A | 27.56 [25.9;29.24] | 3.08 [0.7;5.46] |
| 22B | 24.8 [23.25;26.35] | -2.45 [-4.77;-0.16] |
| 26B | 19.63 [18.05;21.21] | -0.21 [-2.51;2.13] |
| 46B | 17 [15.33;18.64] | 0.43 [-2.11;2.88] |
| 02A | 22.13 [20.57;23.68] | 4.82 [2.54;7.08] |
| 20B | 20.65 [19.05;22.28] | -2.88 [-5.21;-0.56] |
| ***P. fluorescens*** | | |
| 36A | 17.49 [16.14;18.81] | -0.38 [-2.27;1.53] |
| 29A | 17.51 [16.06;18.95] | -1.54 [-3.56;0.45] |
| 39A | 20.35 [19.03;21.66] | -0.71 [-2.6;1.18] |
| 33E | 18.04 [16.74;19.33] | -2.27 [-4.12;-0.37] |
| 03A | 17.76 [16.42;19.1] | -1.97 [-3.92;-0.01] |
| 46A | 20.93 [19.58;22.26] | 0.06 [-1.83;1.98] |

**Table S4.** Model estimates for patch-level fruit set as function of herbivore damage

| Term | Estimate [95% c.i.] |
|---|---|
| $\alpha$ Intercept | 2.6 [2.14;3.06] |
| $\beta$ height (cm) | 0.05 [0.04;0.06] |
| $\beta$ mined leaves (x 10) | -0.06 [-0.1;-0.03] |
| $\sigma$(plot) | 0.35 [0.19;0.49] |
| NB shape $\delta$ | 4.61 [3.11;6.49] |

**Table S5.** Model estimates for impact of hormone treatment on patch-level herbivore abundance in the field (site NP)

| Term | Estimate [95% c.i.] |
|---|---|
| $\alpha$ Intercept (CTR) | -2.31 [-2.92;-1.72] |
| $\beta$ JA | 0.76 [-0.1;1.65] |
| $\beta$ SA | 0.44 [-0.42;1.29] |
| $\sigma$(plot) | 1.31 [1.05;1.63] |
| $\delta$ NB shape | 6.7 [3.98;10.53] |
